# Self-organization of tumor heterogeneity and plasticity

**DOI:** 10.1101/2025.10.09.681487

**Authors:** Carlos Pérez-González, David B. Brückner, Mickael Di-Luoffo, Sophie Richon, Ruchi Goswami, Meryem Baghdadi, Florence Piastra-Facon, Neta Felsenthal, Réda Bouras, Arianna Fumagalli, Mirjam van der Net, Martijn Gloerich, Salvatore Girardo, Jochen Guck, Jacco van Rheenen, Julie Guillermet-Guibert, Edouard Hannezo, Danijela Matic Vignjevic

## Abstract

Phenotypic heterogeneity and plasticity drive tumor growth, metastasis, therapy resistance, and relapse. This heterogeneity is mainly interpreted as a response to external signals from the microenvironment. However, here we show that cancer cells also follow intrinsic self-organized programs that are sufficient to coordinate the spatiotemporal patterning of tumor cell states. By combining quantitative measurements in tumors and organoids with theoretical modeling, we reveal emergent mechanical gradients that orchestrate cell state transitions during colorectal tumor growth. Compression at the tumor center induces a transition from a fetal-like state into a cancer stem cell (CSC) state. The CSC compartment exhibits a characteristic size determined by tumor rheological properties. Once this size is surpassed, a translationally arrested apoptotic core emerges, triggering a shift from homogeneous proliferation to a hierarchical cell turnover. These findings uncover stereotyped programs of self-organization that likely cooperate with the microenvironment to shape tumor heterogeneity and plasticity.

## Introduction

Intratumor heterogeneity represents a major obstacle to the effective treatment of colorectal cancer (CRC) (*1*). Within a tumor, cancer cells adopt distinct phenotypic states that partially mimic the transcriptional and functional heterogeneity of the normal colonic epithelium. For example, proliferative Lgr5+ cancer stem cells (CSC) sustain primary tumor growth by generating short-lived differentiated-like cells (*2–4*). On the other hand, quiescent Mex3a+ (*5*) and/or p27+ (*6*) CSCs do not contribute to tumor growth but can resist chemotherapy. Metastasis and relapse are driven by Lgr5-cells (*7*), which dedifferentiate into Sca1+ fetal-like (*8–11*), L1CAM+ regenerative (*12*) or Emp1+ high-relapse (HRC) (*13*) states. Metastases can also acquire non-canonical neuroendocrine or squamous states (*14*), both associated with poor patient survival. Therefore, intratumor heterogeneity enables diverse cancer cell states to coexist and cooperate, collectively driving tumor growth, metastasis, therapy resistance, and relapse.

Importantly, cancer cells can transition between states in a process known as phenotypic plasticity. For instance, Lgr5-cells that metastasize to the liver or survive therapy can revert to an Lgr5+ state, restoring tumor growth and hierarchy (*2–4, 7, 8*). It is thus crucial to understand which cues induce the acquisition and maintenance of each cancer cell state. During tumor progression, cancer cells accumulate different genetic and epigenetic mutations, but such alterations only explain a small fraction of the tumor phenotypic heterogeneity (*15*). Instead, cell state transitions are mainly interpreted as a response to a microenvironment that is heterogeneous in time and space, providing positional information to the cancer cells (*16–19*). According to this model, dynamic extrinsic signals compose niches that locally induce specific cell states. In contrast, the contribution of tumor-intrinsic mechanisms to establish heterogeneity and plasticity remains largely unexplored.

Here we show that, even in the absence of external positional information, tumors can spontaneously pattern different cell states across time and space guided by emergent tissue mechanical heterogeneities. As tumors grow, cell migration generates tension at the edge, while cell proliferation generates compression at the center. The interplay between these two processes leads to the emergence of a mechanical stress gradient which instructs the radial patterning of Sca1+ fetal-like cells and Lgr5+ CSCs from edge to center. Furthermore, the CSC compartment exhibits a characteristic length, defining a critical size above which the tumor develops a functional hierarchy characterized by a decreased biosynthetic capacity and the development of an apoptotic core. Together, these findings reveal intrinsic feedback loops between force, proliferation and fate that govern spatiotemporal dynamics of tumor plasticity, pointing towards self-organization as a driver of CRC heterogeneity and plasticity.

### Growing tumor organoids recapitulate the spatiotemporal patterning of Sca1+ and Lgr5+ cancer cells observed in tumors

To study the spatiotemporal dynamics of cell states in CRC, we implanted mouse tumor organoids (APC^-/-^, KRAS^G12D^, p53*-/-*) in the interscapular fat pad of immunodeficient mice and collected samples at different timepoints (Fig. 1A). These organoids constitutively express RFP to visualize tumor morphology, and a fluorescent reporter of Lgr5 (Lgr5:EGFP-DTR(*3*)) to label CSCs. Additionally, we performed immunostainings of Sca1 as a marker of a fetal-like state. Small tumors (2-7 days post-injection) mostly contained Sca1+ Lgr5-cells, with few Lgr5+ cells dispersed within the tumor. As tumors grew (1-2 weeks post-injection), Sca1 became restricted to the tumor edge, and Lgr5 increased at the tumor center. Larger tumors (3 weeks) maintained this radial organization—Sca1 at the periphery and Lgr5 toward the center—but the tumor core developed a more complex cell-state distribution, with alternating Sca1+Lgr5− and Sca1−Lgr5+ zones in a seemingly periodic pattern. This shows that cell states become stereotypically patterned as tumors grow, defining at least three compartments from tumor edge to core (Fig. 1A, Fig. S1A-B).

**Fig. 1.**
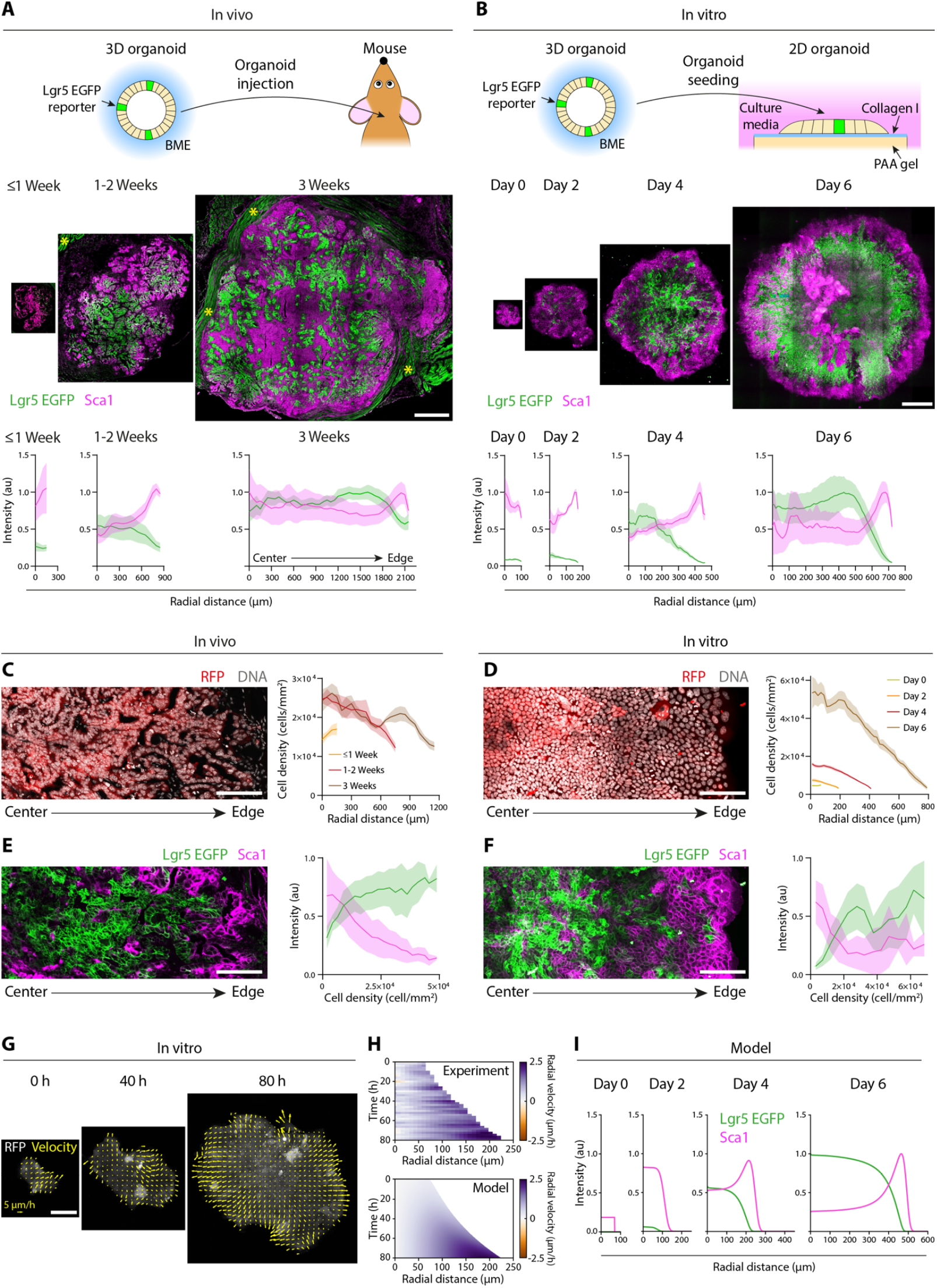
Emergent cell density gradients coordinate the spatiotemporal patterning of tumor cell states. **(A-B)** Top: scheme of the *in vivo* (A) and *in vitro* (B) experimental setups. Middle: Lgr5 EGFP and immunostaining of Sca1 in subcutaneous tumor sections (A) and expanding organoids (B) at different timepoints. Scalebar = 1000 μm (A) and 300 μm (B). Yellow asterisk labels autofluorescent muscle, detected by its RFP-negative signal (see Fig. S1A). Bottom: average radial intensity profiles of Lgr5 EGFP and Sca1 intensity. Mean ± SEM. For tumors (A), N= 6 (≤1 week), 6 (2 weeks) and 5 (3 weeks). For organoids (B), n= 9 (Day 0), 4 (Day 2), 6 (Day 4) and 5 (Day 6) organoids from N=2 (Day 0-4) and N=3 (Day 6) independent experiments. **(C-D)** Left: cancer cells (RFP) and DNA (DAPI) in 3 weeks tumors (C) and 4 days organoids (D). Scalebars = 100 μm. Right: radial cancer cell density profiles in tumors (C) and organoids (D). Mean ± SEM. Tumors are the same as in A-B. **(E-F)** Left: Lgr5 EGFP and immunostaining of Sca1 in 3 weeks tumors (C) and 4 days organoids (D). Scalebars = 100 μm. Right: cell state intensity as a function of local cell density in tumors (C) and organoids (D). Mean ± SD. For tumors (E), n = 2612 local measurements from the 17 tumors in A. For organoids (F), n = 1007 local measurements from the 24 organoids in B. (**G)** Velocity maps of an expanding organoid. Scalebar = 100 μm. **(H)** Average experimental (top) and model (bottom) kymograph of radial velocity in expanding organoids. Top: n=10 organoids from N=3 independent experiments. **(I)** Model simulations of Lgr5 and Sca1 radial profiles from day 0 to day 6.

To dissect the mechanisms driving spatiotemporal patterning of cell states, we studied organoid growth in a controlled biochemical and mechanical *in vitro* environment. Specifically, we limited external biochemical signals by using a minimal culture medium devoid of classical niche factors (Wnt, R-Spondin, Epidermal Growth Factor - EGF- or Noggin). Given that tumors maintain a glandular architecture and apicobasal polarity, we cultured organoids as 2-dimensional monolayers (*20–22*) on soft substrates (5kPa Young’s Modulus) coated with collagen I (Fig. 1B). Furthermore, initial monolayer geometry was controlled by patterning organoids into 200 μm circular stencils, which were removed upon cell attachment to allow tissue expansion (*23*). At day 0, organoids systematically contained Sca1+ Lgr5-cells, showing that our seeding strategy either selects or induces a fetal-like state. At days 2 and 4, cells at the organoid center lost Sca1 and increased Lgr5 expression. By day 6, however, Lgr5 was downregulated at the center and restricted to a rim adjacent to the peripheral Sca1+ cells. At this point, Sca1 became re-expressed in some central regions. Growing organoids thus largely recapitulate the spatiotemporal patterning of cell states observed *in vivo* (Fig. 1B, Sup. fig. 1C-D).

These observations are particularly striking, as tumor radial patterning has traditionally been attributed to the limited diffusion of microenvironmental growth factors, nutrients and oxygen from edge to center (*24, 25*). Yet, our *in vitro* system lacks stromal-derived growth factors, and its 2D geometry limits the formation of diffusive gradients compared to 3D tissues. To confirm this and rule out other modes of gradient formation, e.g. at the cell-substrate interface as seen in embryonic cell monolayers (*26–28*), we cultured organoids on permeable filters where media was equally accessible at the basal and apical monolayer surface. Even under these highly uniform external conditions, Lgr5 expression became radially patterned (Fig. S2). Our results thus reveal the existence of a tissue-intrinsic mechanism controlling the spatiotemporal patterning of cell states.

### An interplay between migration and proliferation leads to the emergence of a density gradient that correlates with cell states

Given the apparent absence of external biochemical gradients, we explored the possibility of mechanical gradients as organizers of tumor patterning. Small organoids and tumors exhibited a homogeneously low cell density but, as they grew, developed a radial gradient of increasing density from edge to center (Fig. 1C-D). Local cell density correlated with cell states for all timepoints, with Sca1+ cells systematically located in low density regions, and Lgr5+ cells in high density regions (Fig. 1E-F). Importantly, this state-density relation was conserved in mouse tumors growing at the colon and liver (Fig. S3), highlighting its independence from the microenvironment and strongly suggesting an intrinsic nature. We thus hypothesized that spatiotemporal changes in cell density during tumor growth could guide the dynamics of cell state transitions.

To quantitatively test this hypothesis, we developed a theoretical framework to predict the density, stress and velocity fields in the tumor. As we focus on long timescales compared to the dynamics of cell-cell rearrangements, the tissue can be described as an active viscous fluid (SI Theory) (*29, 30*). Experimental measurements revealed that cell proliferation followed non-monotonic patterns during organoid expansion: it was homogeneous at the beginning of the experiment, increased at the tissue center as cells expressed Lgr5, and became progressively restricted to the edge as the tissue center became crowded (Fig. S4). A simple logistic growth model quantitatively fitted the measured growth rates, allowing us to extract all proliferation parameters (SI Theory). For force balance, we assume frictional contact with the substrate which generically predicts a build-up of density and velocity gradients during growth and expansion (*31–33*). Consistent with our model, live-imaging and particle-image velocimetry (PIV) analysis of spreading organoids revealed a radial velocity gradient increasing linearly from center to edge (Fig. 1G-H, Movie S1). The interplay between velocity and proliferation naturally explains the establishment of a radial cell density gradient and quantitatively fits the expected behavior of an actively expanding fluid. By imposing the experimental state-density relation, our model recapitulates the radial patterning of Sca1+ cells at the edge and Lgr5+ at the center as the tumor grows without the need for external positional information (Fig. 1I).

### Growth-induced mechanical compression triggers an Lgr5 CSC state

The active fluid model provided a key prediction: tissue confinement should speed up the appearance of Lgr5+ CSCs (SI Theory). This arises because confinement limits outward cell movement, steepening the radial cell density gradient as cells proliferate. To test this prediction experimentally, we confined organoids into 200 μm diameter circular collagen I micropatterns surrounded by a non-adhesive surface (*34*). Live-imaging revealed that confinement triggers an earlier onset and a faster increase in Lgr5 expression compared to spreading organoids, despite the identical exposure to culture media and substrate mechanochemical properties (Fig. 2A-D, Fig. S5D, Movie S2).

**Fig. 2.**
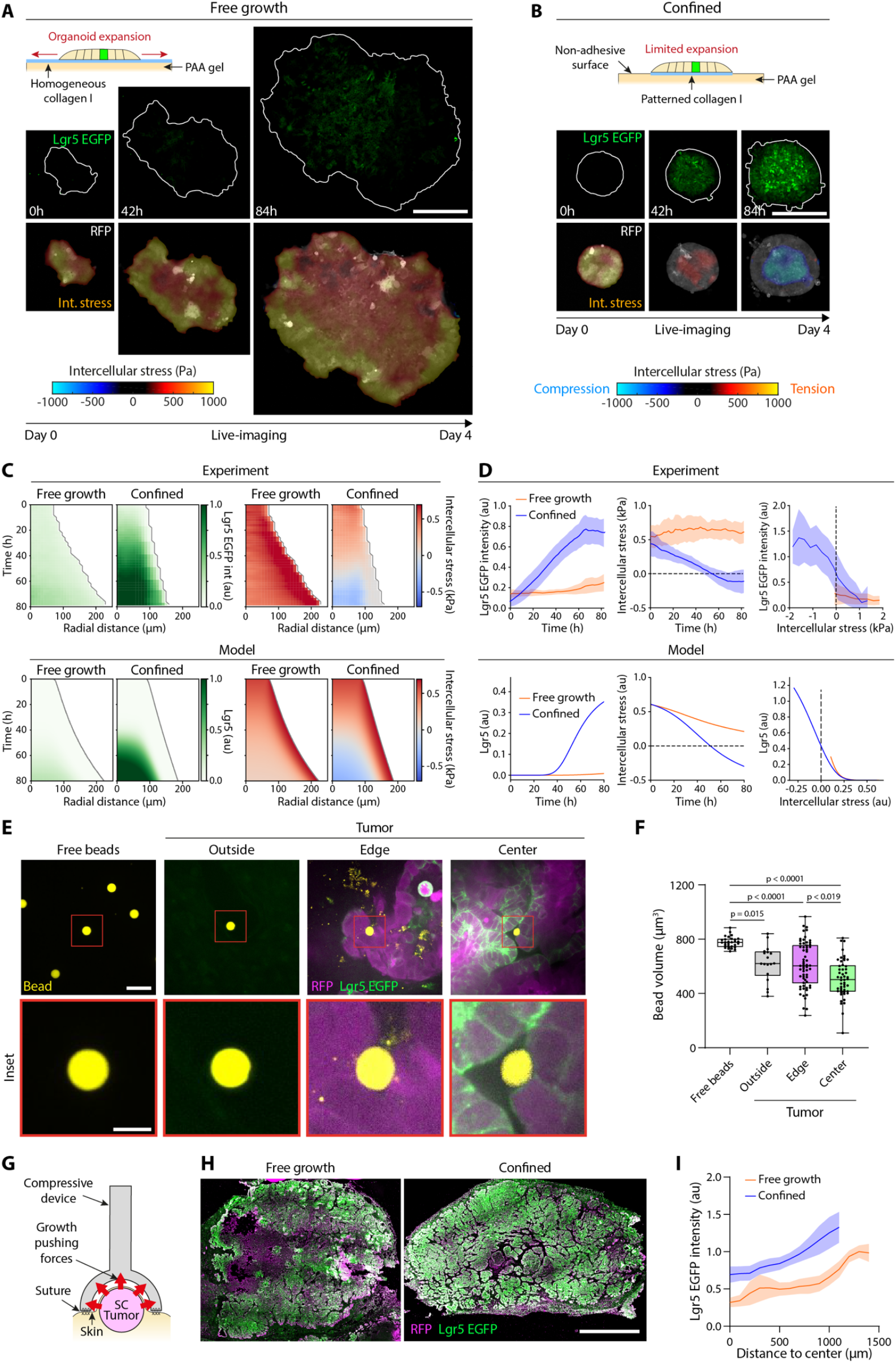
Mechanical compression induces an Lgr5 CSC state. **(A-B)** Live-imaging of Lgr5 EGFP (top) and intercellular stress maps overlayed on RFP signal (bottom) for freely expanding (A) and confined (B) organoids. Scalebar = 200 μm. **(C)** Average experimental (top) and model (bottom) kymographs of Lgr5 and intercellular stress for confined and freely growing organoids. n=10 (free growth) and 22 (confined) organoids from N=3 (free growth) and 4 (confined) independent experiments. **(D)** Left and center: Time evolution of average Lgr5 intensity (left) and intercellular stress (center) for experiment (top) and model (bottom). Mean ± SD. Same organoids as in C. Right: Local Lgr5 intensity as a function of intercellular stress for experiments (top) and model (bottom). Mean ± SD of n = 615025 (free growth) and 702166 (confined) local measurements from all organoids and timepoints in C. **(E)** Representative examples (top) and insets (bottom, red squares) of polyacrylamide beads integrated at different locations of a 1-week tumor. Scalebar = 30 μm (top) and 10 μm (bottom, inset). **(F)** Bead volume at different locations of 1-week tumors. Points indicate single beads, boxes indicate the mean, quartiles and minimum to maximum range. n = 30 (free beads), 18 (outside), 63 (edge), 50 (center) from N=5 independent tumors. One-way ANOVA test with multiple comparisons. **(G)** Scheme of the compression device. **(H)** Immunostaining of Lgr5 EGFP and RFP in freely growing (left) and confined (right) tumors. **(I)** Radial profiles of Lgr5 EGFP intensity in freely growing and confined tumors. Mean ± SEM of N=5 (free growth) and 6 (confined) tumors.

The model further predicted that the transition from fetal-like to a CSC state in spreading vs confined organoids should be accompanied by a change in the mechanical state of the tissue: while expanding tissues are predicted to exhibit tensile stress profiles, confined tissues are expected to eventually switch to compressive stresses, since proliferation-induced pushing is no longer balanced by outwards expansion. To test this, we measured cell-substrate forces using Traction Force Microscopy (TFM) (*35*), and intercellular stresses using Monolayer Stress Microscopy (MSM) (*36*) (Fig. 2A-D, Fig. S5, Movie S2,3). Consistent with the model, both spreading and confined organoids were initially under tension but evolved differently. Spreading organoids exhibited a constant inwards-pointing traction peak at the tissue edge and an overall tensile stress profile. Intercellular stresses were high at the edge and decayed towards the tissue center. Confined organoids, instead, experienced a progressive decrease in tissue tension, which became compressive after 60h. Interestingly, by plotting the intercellular stress as a function of Lgr5 EGFP intensity (Fig. 2D), we observed that Lgr5 is specifically upregulated in mechanically compressed regions, suggesting that proliferation-driven compressive forces guide the transition from Sca1 into the Lgr5 state.

To test whether mechanical compression could induce the Lgr5 CSC state also *in vivo*, we first performed cell-scale stress measurements in tumors using polyacrylamide beads as force sensors (5.9±2 kPa Young’s Modulus, 14.9 ± 0.5 μm diameter) (*37–39*). We co-injected organoids and beads in the interscapular fat pad of mice and harvested the tumors for 1 or 2 weeks. Tumors were then extracted, fixed and cleared for high-resolution 3D imaging. Polyacrylamide beads integrated in different regions of the tumor (Fig. S6A), allowing for the comparison of local stresses outside the tumor, at the tumor edge, and the tumor center. Unfortunately, bead fixation and clearing alone triggered a decrease in bead volume, hindering the calculation of absolute stress values (Fig. S6B). Yet, both in 1- and 2-week tumors, beads in all tumor regions exhibited an average decrease in volume compared to fixed and cleared beads, indicating that tumors grow under compressive stress (Fig. 2F, Fig. S6C-D). Interestingly, beads at the tumor center had a significantly smaller volume and a higher eccentricity than beads at the tumor edge (Fig. 2F, Fig. S6C-D). These measurements reveal the existence of an *in vivo* compressive stress gradient that may induce the appearance of Lgr5+ CSCs at the tumor center.

To directly test the functional role of compressive forces *in vivo*, we implanted a compressive device around 100 mm^3^ subcutaneous tumors (Fig. 2G, Fig. S7A-B)(*40*). The device confined tumors for 4 days, which was sufficient to observe a significant reduction in tumor growth (Fig. S7C-D). Tumor harvesting and staining revealed an increased Lgr5 EGFP fluorescence intensity in confined tumors compared to those freely expanding (Fig. 2H-I). This result shows that, even when cancer cells are exposed to a complex *in vivo* microenvironment, growth-induced compression promotes an Lgr5 CSC state.

### Tissue hierarchy emerges above a critical tissue size

Having shown that growth-induced compression can induce a transition from Sca1+ fetal-like to Lgr5+ CSC state, we investigated the origin of the third compartment - characterized by a decreased Lgr5 expression and the reappearance of Sca1 in the center - observed in 3 weeks tumors and 6 days spreading organoids (Fig. 1A-B).

*In vivo*, the appearance of this compartment correlated with the compartmentalization of cell proliferation at the tumor edge and the establishment of an apoptotic core (Fig. 3A-B). This patterning has been classically attributed to hypoxia (*24, 25*), where limited oxygen diffusion at the core leads to an inhibition of proliferation and subsequent cell death. Yet, a deeper analysis of the tumor apoptotic core revealed the presence of ∽100 μm stripes of proliferative Lgr5+ cells adjacent to apoptotic Sca1+ cells (Fig. 3C-D, Fig. S8). Lgr5+ cells look viable based on their morphology (RFP), suggesting an adaptation to the environment even if local hypoxia could exist. 2D autocorrelation of Lgr5 EGFP intensity in central tumor areas reveals a spatial periodicity of ∽200 μm (Fig. S8B). The periodicity and characteristic size of the Lgr5 stripes are difficult to explain exclusively by monotonic radial oxygen gradients, suggesting a contribution of tumor-intrinsic mechanisms to the establishment of such patterning. The spatial organization of Lgr5, proliferation and death within each stripe is reminiscent of the crypt-plateau organization of the homeostatic colonic epithelium (*41*). It is therefore appealing to think that these Lgr5 stripes compose hierarchical units undergoing a constant cell turnover through proliferation and death.

**Fig. 3.**
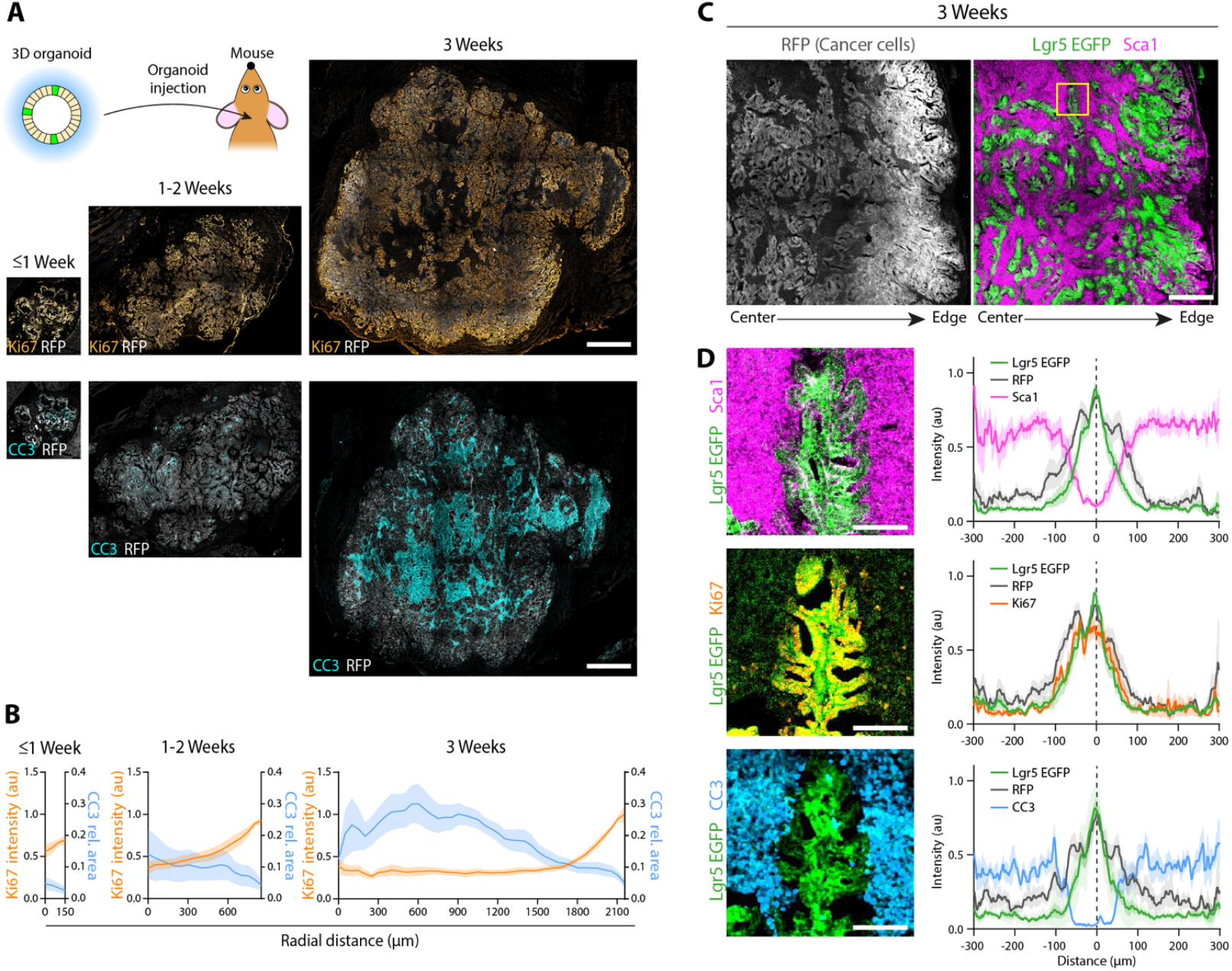
Periodic hierarchical units develop at the apoptotic core of large tumors. **(A)** Immunostainings of Ki67 and cleaved-caspase 3 (CC3) in subcutaneous tumors expressing RFP at different timepoints after injection. Tumors are the same as shown in Fig. 1A, stained in consecutive sections. Scalebar = 1000 μm. **(B)** Average radial intensity profiles of Ki67 intensity and CC3 tumor area coverage. Mean ± SEM for N= 6 (≤1 week), 6 (2 weeks) and 5 (3 weeks) tumors. Tumors are the same as shown in Fig. 1A, stained in consecutive sections. **(C)** Representative image of a 3-weeks subcutaneous tumor expressing RFP (left) and Lgr5 EGFP stained for Sca1 (right). Lgr5 stripes are located at the tumor center. Yellow square indicates the striped shown in D. Scalebar = 500 μm. **(D)** Left: representative image of an Lgr5 stripe at the tumor core stained for Sca1 (top), Ki67 (middle) and cleaved-caspase 3 (CC3, bottom). Images show the same stripe in consecutive sections. Right: average linescans of Lgr5 EGFP, RFP, Sca1 (top), Ki67 (middle) and CC3 (bottom) intensity as a function of the distance to the Lgr5 central maximum (distance =0). Mean± SEM of n=9 stripes from N=3 tumors. Scalebar = 100 μm.

We then used organoids to study the intrinsic mechanisms and functional consequences of the decrease in Lgr5 at the tumor center. We noted that the central Lgr5-compartment was absent in 200 μm confined organoids (Fig. 2B), suggesting that, as in tumors, such patterning only appears above a critical tissue size. To dissect the role of organoid size in cell state patterning, we confined organoids into circular collagen I patches of increasing diameters: 200 μm, 500 μm, 1000 μm and 2000 μm (Fig. 4A). At day 4, as shown above, 200 μm organoids exhibited uniformly high Lgr5 EGFP intensity. In contrast, in larger organoids, Lgr5 was restricted to a peripheral rim of ∽200 μm, with a progressive decrease toward the center (Fig. 4A). Remarkably, the width of this rim remained constant regardless of organoid size (Fig. 4A, Fig. S9A).

**Fig. 4.**
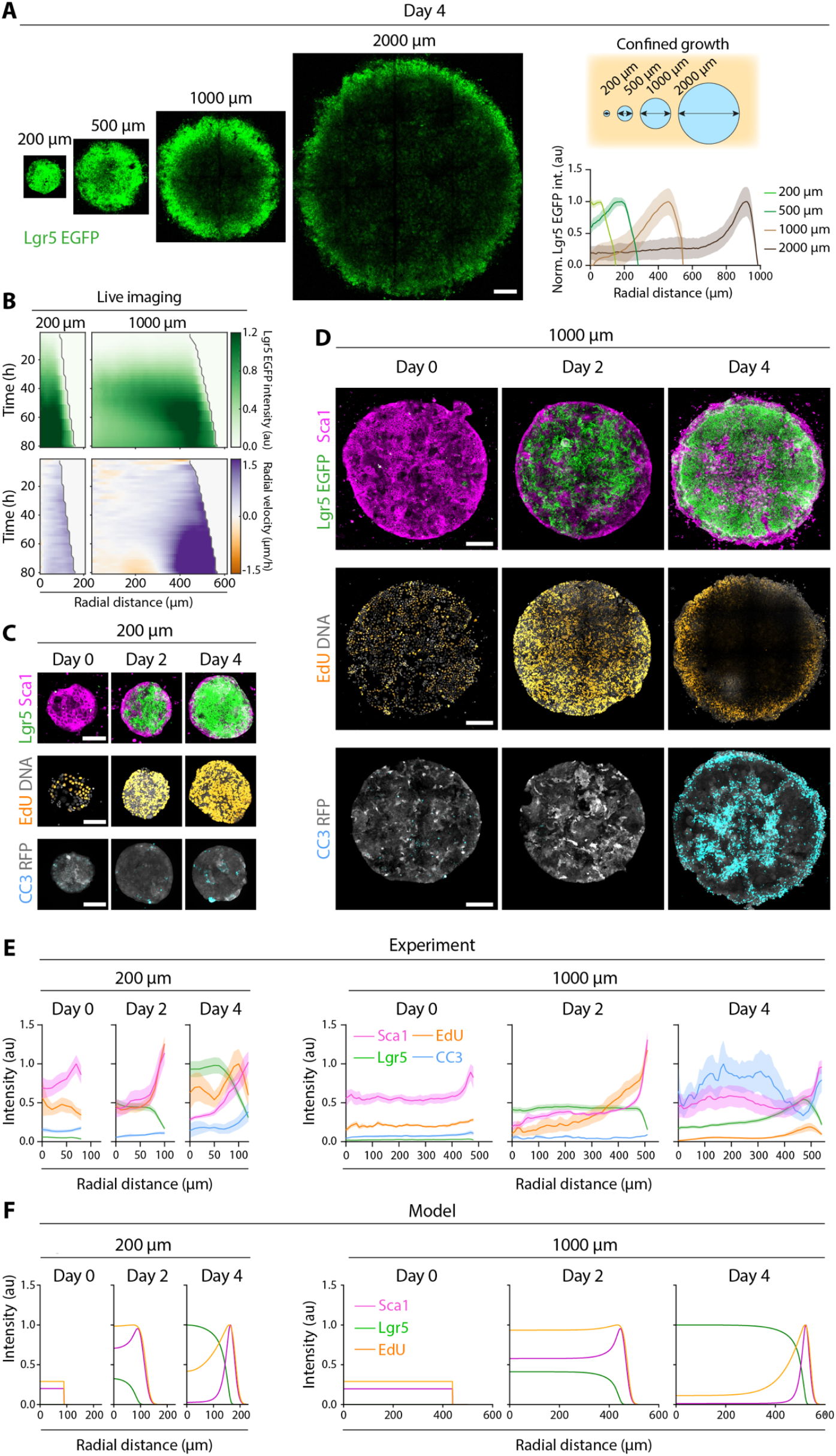
Functional hierarchy emerges above a critical tissue size defined by the characteristic length of the CSC niche. **(A)** Left: Live-imaging of Lgr5 EGFP in patterned organoids of different sizes at day 4. Scalebar = 200 μm. Right: Normalized Lgr5 EGFP radial profiles for all sizes. Mean ± SEM of n = 26 (200 μm), 13 (500 μm), 15 (1000 μm) and 3 (2000 μm) organoids from N = 4 (200 μm, 500 μm, 1000 μm) and 3 (2000 μm) independent experiments. Normalization was performed to better visualize the constant Lgr5 EGFP peak width. **(B)** Average kymographs of Lgr5 EGFP intensity and radial velocities for 200 μm and 1000 μm organoids. Representative images are shown in Fig. S9 B-C. n=22 (200 μm) and 12 (1000 μm) organoids from N=3 independent experiments. **(C-D)** Immunostaining of Sca1, EdU and cleaved-caspase 3 (CC3) in 200 μm (C) and 1000 μm (D) organoids at Day 0, 2 and 4. Note that each staining is performed in an independent organoid. Scalebar = 100 μm (C) and 200 μm (D). **(E-F)** Time evolution of radial profiles of Lgr5, Sca1, Proliferation/EdU and CC3 in experiments (E) and model simulations (F). Mean ± SEM. 200 μm: Sca1, n= 9 (day 0), 5 (day 2) and 3 (day 4) organoids from N=2 experiments; EdU, n= 15 (day 0), 26 (day 2) and 16 (day 4) organoids from N=5 experiments; CC3, n= 8 (day 0), 9 (day 2) and 10 (day 4) organoids from N=3 experiments. For 1000 μm: n= 6 (day 0), 4 (day 2) and 5 (day 4) organoids from N=2 experiments; EdU, n= 10 (day 0), 17 (day 2) and 10 (day 4) organoids from N=5 experiments; CC3, n= 6 (day 0), 5 (day 2) and 6 (day 4) organoids from N=3 experiments.

To investigate the development of the Lgr5 rim, we performed live imaging of 200 μm (small) and 1000 μm (large) organoids (Fig. 4B, Fig. S9B-C, Movie S4). In both sizes, Lgr5 expression homogeneously increased during the first 2 days. However, while small organoids maintained a high and homogeneous Lgr5 EGFP intensity by day 4, large organoids exhibited a sudden decrease in Lgr5 EGFP intensity at the center, leading to the establishment of a ∽200 μm Lgr5 peripheral rim. PIV analysis revealed that cells generally moved outwards, reflecting the slow tissue expansion. However, the appearance of the Lgr5 rim was paralleled by a switch in cell velocity, with cells moving from the Lgr5+ pool at the edge towards the organoid center (Fig. 4B, Fig. S9B-C, Movie S5).

To functionally characterize the differences between small and large organoids, we stained for Sca1, proliferation (EdU) and apoptosis (cleaved-caspase 3) at day 0, 2 and 4 (Fig. 4C-E, Fig. S9D-E). Both small and large organoids had a uniformly high Sca1 expression at day 0 but transitioned towards Lgr5 by day 2, restricting Sca1 to the outer-most cells of the organoid edge. Only in large organoids, however, some cells at the tissue center lost Lgr5 and re-expressed Sca1 by day 4 (Fig. 4C-E). Small organoids proliferated throughout the 4 days and exhibited minimal apoptosis, with the few dying cells preferentially located at the tissue edge, probably due to the organoid growth against a boundary. Large organoids, instead, progressively restricted cell proliferation to the Lgr5 peripheral rim and developed an apoptotic center by day 4 (Fig. 4C-E, Fig. S9D-E). Altogether, this shows that small organoids mostly contain Lgr5 CSCs in a proliferative state. In contrast, large organoids establish a functional hierarchy whereby Lgr5 CSCs proliferate at the edge, move towards the center while losing stemness, and die. This striking patterning dynamics and its reproducibility suggest that cancer cells spontaneously develop a functional hierarchy upon reaching a critical tissue size defined by the characteristic length of the CSC compartment.

### Single cell RNA sequencing reveals the time evolution of cell states and a decrease in translation in large organoids

Interestingly, our computational model successfully predicted the constant length-scale of Sca1 patterning in 200 vs 1000 μm patterns, and the decrease of cell proliferation in the center of large patterns (Fig. 4F). However, it failed to predict the emergence of the third compartment in large organoids at day 4 (Fig. 4F): in the simple picture in which Lgr5 is positively regulated by cell density, no downregulation of Lgr5 in the tissue center is expected as this is where cell density is maximal. This suggests that, while Lgr5 is initially upregulated by cell density, an additional factor may cause downregulation at very high densities. In search of such a mechanism, we performed single cell RNA sequencing of small (200μm) and large (1000 μm) organoids at days 0, 2 and 4. During this time, cells adopted different phenotypic states represented by 7 clusters (Fig. S10, Fig. 5A). Based on the expression of known genes and signatures from the intestinal epithelium (*13, 42–49*), we labeled each cluster as: cells under Endoplasmatic Reticulum (ER) stress, fetal-like cells, two clusters of proliferative cancer stem cells (proCSC1, proCSC2), two clusters of Lgr5+ CSCs (CSC1 and CSC2) and differentiated-like cells in the secretory and absorptive lineages (Fig. 5A,C, Fig. S11C, Fig. S12). The time evolution of these cell states reveals that organoids at day 0 contain cells in an ER Stress state and in a fetal-like (Ly6a, Anxa1) and/or high-relapse cell (HRC) states (Emp1) (Fig. 5B,D, Fig. S11A-C). Interestingly, these cells also exhibit a high Yap signaling score, a well-known mechanosensor (*50, 51*) and regulator of the fetal-like program (*8, 43, 52*). At day 2, cells transition into a proliferative CSC state, characterized by the expression of proliferation markers (mKi67, Pcna) and the abundance of cells in the S/G2M phases of the cell cycle. Already by day 2, some cells adopted a differentiated-like absorptive state. By day 4, proliferation decreased, and cells adopted a CSC state positive for markers of adult intestinal stem cells and Wnt signaling (Lgr5, Ephb2, Lrig1, Ascl2). Interestingly, many CSCs from the CSC1 group adopted a slow-proliferative signature, characterized by the Mex3a state and the presence of cells in a G1 phase of the cell cycle (Fig. 5B,D, Fig. S11A-D). The rest of the cells adopted differentiated-like states either in the secretory (Muc2, Reg4, Atoh1) or absorptive (Krt20, Aldob, Fabp2) lineages (Fig. 5B,D, Fig. S11A-D). This data validates the use of Sca1 and Lgr5 as markers of fetal-like and CSC states.

**Fig. 5.**
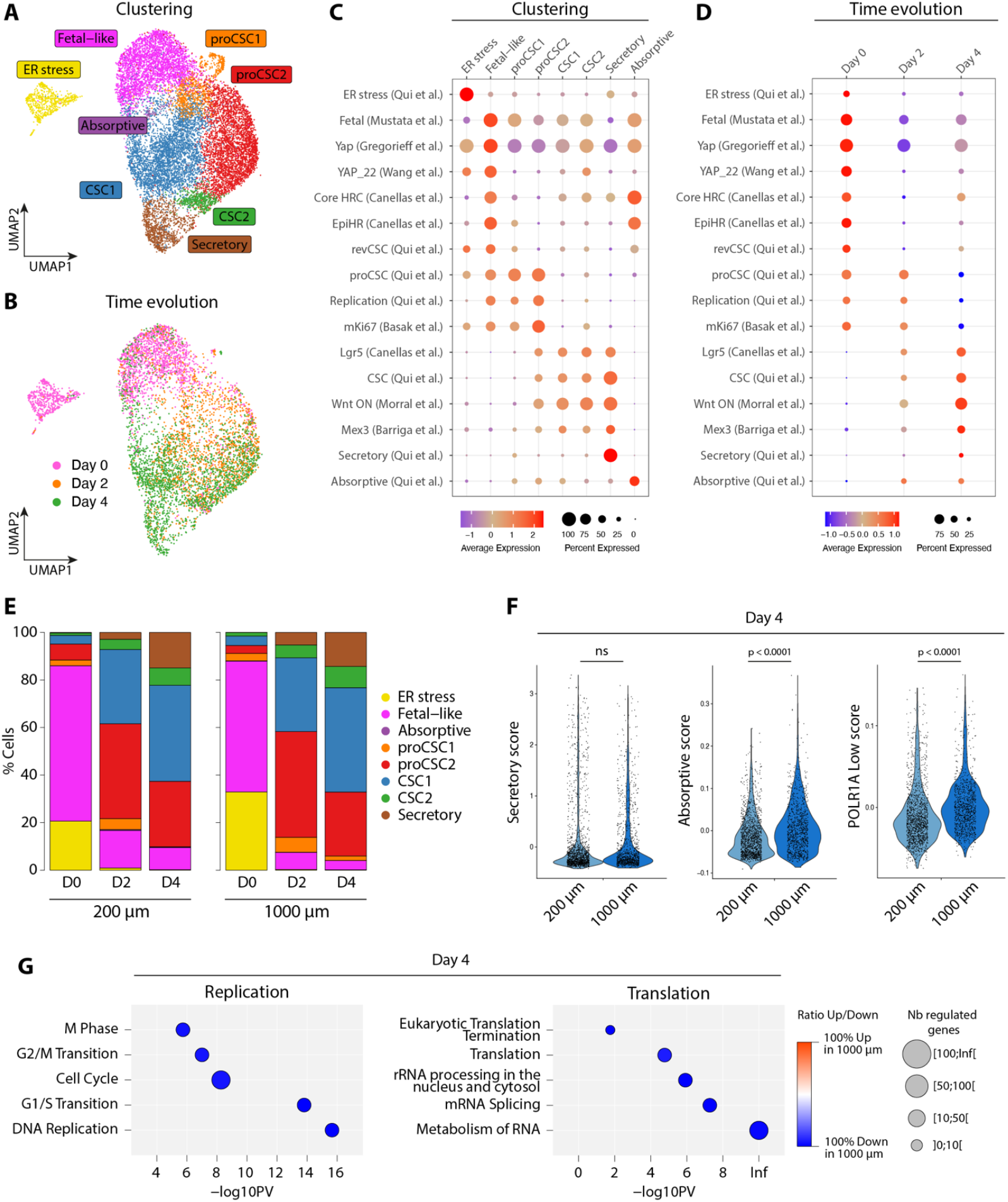
Single cell RNA sequencing reveals cell state trajectories of proliferative and hierarchical organoids. **(A-B)** UMAP showing the cluster classification (A) and time after seeding (B) of all sequenced single cells from small (200 μm) and large (1000 μm) organoids. **(C-D)** Score of published signatures per cluster (C) and time after seeding (D). **(E)** Cluster proportion for small (left) and large (right) organoids at day 0, 2 and 4 after seeding. **(F)** Violin plot comparing the score of secretory (left), absorptive (center) and POLR1A Low (right) signatures between small and large organoids at day 4. T-test was performed to assess significance. **(G)** Differential expression analysis between small (200 μm) and large (1000 μm) organoids at day 4. REACTOME terms related to replication (left) and RNA metabolism and translation (right) are shown.

Surprisingly, except for an increase in the absorptive score in large organoids at day 4 (Fig. 5F), we did not observe different trajectories between small and large organoids based in the time evolution of cluster proportions (Fig. 5E). We thus performed differential expression analysis at day 4 to detect transcriptional changes beyond our cell-state classification, and interpreted these changes using annotation databases (KEGG, Reactome and Gene Ontology). For all databases, we observed a decreased expression of genes related to the cell cycle in large organoids (Fig. 5G, Fig. S13A), which was consistent with our experimental observations (Fig. 4C-E). Surprisingly, large organoids also exhibited a systematic downregulation of ribosomal RNA synthesis, RNA processing and translation (Fig. 5G, Fig. S13B). Cancer cell biosynthetic capacity has been associated with stemness and is radially patterned in tumors, where high biosynthetic cells are found at the tumor edge (*46*). Signatures of biosynthetic capacity (POLR1A high and POLR1A low) showed that organoids at day 0 and 2 were highly biosynthetic, whereas at day 4 they switched into a low biosynthetic state (Fig. S11E). This low biosynthetic state was significantly higher in large organoids (Fig. 5F). Together, these findings led to the hypothesis that the formation of the third compartment is driven by a decrease of translation.

### Translational arrest leads to a functional loss of stemness

To experimentally validate the decrease in translation, we incubated small and large organoids at day 4 with the amino acid analog O-propargyl-puromycin (OPP). Quantifying OPP incorporation by its fluorescent intensity provided spatial profiles of cell biosynthesis, which were strikingly similar to the intensity profiles of Lgr5 EGFP (Fig. 6A). Small organoids exhibited a plateau in OPP intensity at the center, whereas large organoids exhibited an OPP peak at the Lgr5 EGFP rim near the edge that decayed towards the center. Moreover, live-imaging revealed that, as Lgr5 EGFP decreases in the center of large organoids, a similar radial gradient of RFP intensity emerges, even though RFP is constitutively expressed by all cancer cells (Fig. 6B, Movie S6). Even tumors developed an RFP gradient from edge to center by week 3 (Fig. 6B). This suggests that, at least partially, the decrease in Lgr5 EGFP intensity at the tissue center may be due to a reduction in translation, and not necessarily due to changes in Lgr5 transcription. Consistently, normalizing Lgr5 EGFP intensity by the RFP intensity largely compensated for the differences between small and large organoids (Fig. 6C-E).

**Fig. 6.**
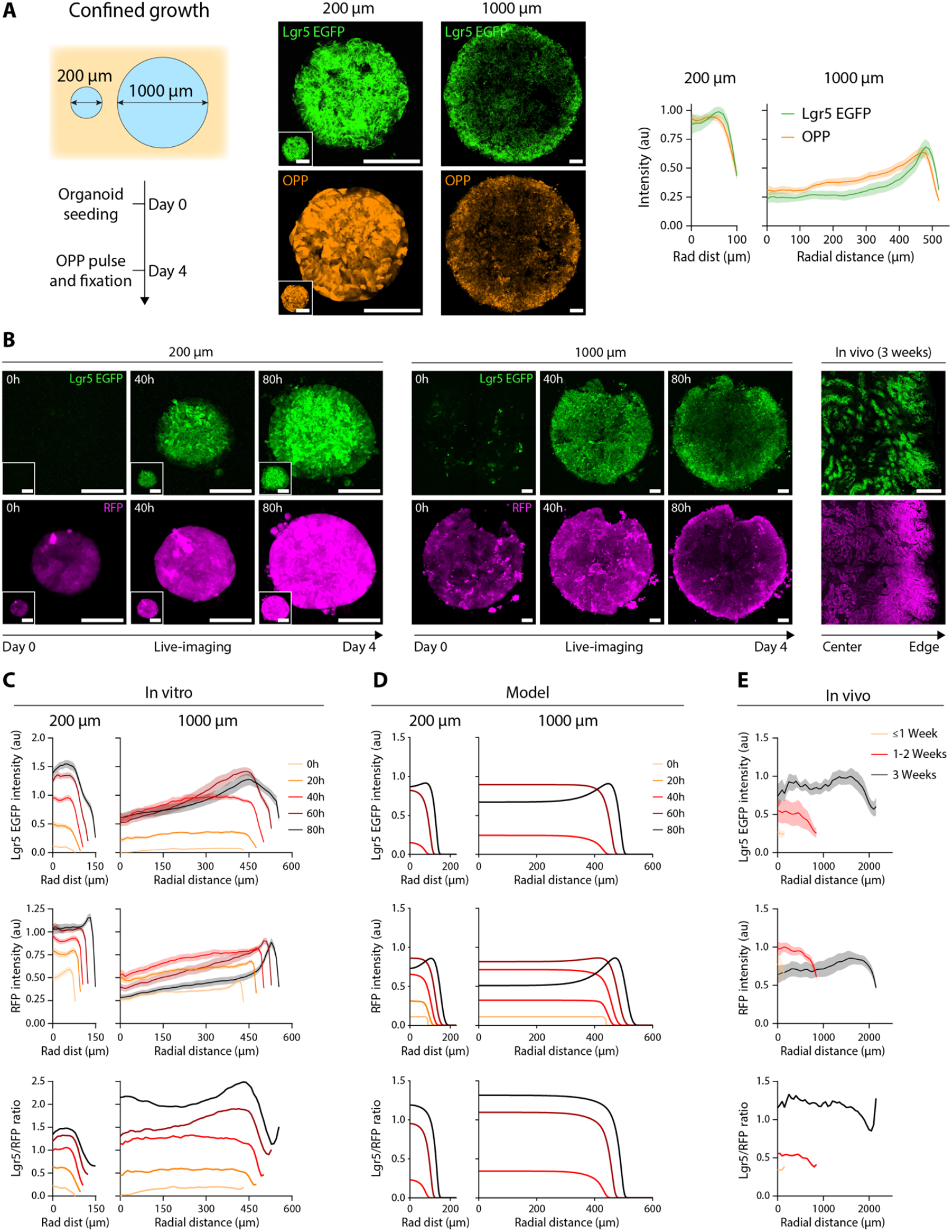
Biosynthetic arrest explains the decay of Lgr5 at the center of hierarchical organoids and tumors. **(A)** Left: Scheme of the OPP pulse experiment. Center: imaging of Lgr5 EGFP and OPP incorporation in small (200 μm) and large (1000 μm) organoids. Scalebar = 100 μm (Small organoids are scaled as large organoids in the bottom left white squares). Right: radial intensity profiles of Lgr5 EGFP and OPP for both organoid sizes. Mean ± SEM of n=19 (small) and 8 (large) organoids from N=3 experiments. **(B)** Representative Lgr5 EGFP and RFP images in small organoids (left), large organoids (center) and 3 week tumors (right). Organoids are imaged live. Scalebar = 100 μm (Small organoids are scaled as large organoids in the bottom left white squares). **(D-F)** Time evolution of radial profiles of Lgr5 EGFP (top), RFP (middle) and Lgr5/RFP ratio (bottom) in organoids (D), model simulations (E), and tumors (F). Organoids are the same as in Fig. 3B. Tumors are the same as in Fig. 1A.

Our observations, together with observations in yeast showing that compressive stress limits both proliferation and translation through macromolecular crowding (*53*), suggests that enhanced compression in central regions cause the decay in translation. To test this scenario, we amended a density-dependent translation inhibition in our computational model, which allowed to capture the observed Lgr5 profiles. The emergence of the third compartment in the center of the tissue was caused by the increasing buildup of cell density accompanied by compressive stresses and a decreased translation of Lgr5 and RFP (Fig. 6D). Indeed, in the experiments, the center of large organoids by day 4 becomes highly crowded and undergoes folding around the time when Lgr5 and RFP become downregulated, whereas in small organoids, folding is less pronounced (Fig. S14, Movie S7). This is consistent with model predictions of an increased compression at the center of 1000 μm organoids (SI Theory).

## Discussion

Tumor cell state patterning is primarily attributed to extrinsic microenvironmental signals. Indeed, numerous stromal-derived factors induce specific cell phenotypic states (*16, 17, 19, 42*). Moreover, the limited diffusion of these factors -along with oxygen and nutrients-can account for the radial patterning of cell states within tumors. However, the development of organoids suggests the existence of additional intrinsic mechanisms controlling tumor heterogeneity and plasticity (*54, 55*). Patient-derived organoids greatly recapitulate the heterogeneity of their tumor of origin, despite being cultured in a homogeneous environment lacking positional cues (*14, 56*). Furthermore, as tumors acquire mutations, cancer cells progressively become independent of external signals (*42*), supporting the role of cell-intrinsic cues as inducers of specific cell states. Yet, in a genetically homogeneous population, intrinsic cues at the single cell level cannot explain the emergence of heterogeneity at the tissue level. Our data demonstrates that, as tumors grow, naturally emerging heterogeneities coordinate the spatiotemporal patterning of cell states, uncovering stereotyped tissue-intrinsic programs as organizers of intratumor heterogeneity. Interestingly, these programs mirror some of the transitions observed during liver metastatic growth, where Emp1+ HRC and/or Sca1+ fetal-like cells become Lgr5+ CSCs and establish a functional hierarchy (*7, 8, 13*).

Mechanical signals are increasingly recognized as cell fate regulators. In the healthy intestinal epithelium, substrate stiffness (*20, 57–59*), viscoelasticity (*60*), stretch (*59, 61*), shear stress (*62*), hydraulic pressure (*63, 64*) or tissue curvature (*65, 66*) govern the balance between stemness and differentiation. In colorectal cancer, integrin-mediated traction forces promote a fetal-like state (*52*), and adhesion by hemidesmosomes promotes a chemoresistant Lgr5+ p27+ state (*6*). Moreover, alterations in extracellular matrix and cell mechanical properties are essential for reprogramming normal cells into tumor precursors and sustaining tumor growth (*67–69*). In this study, we show that the dynamic interplay between cell proliferation and migration during tumor growth generates spatial gradients of mechanical stress, giving rise to distinct mechanical niches that organize cell states into defined tumor compartments. These compartments exhibit a characteristic hydrodynamic length-scale determined by tissue rheological properties, which governs the sequential emergence of cell states as tumors grow. Initially homogeneous and fetal-like, tumors first establish a central Lgr5 CSC domain, which above a critical size, develops a translationally arrested apoptotic core. This transition has profound implications for tumor architecture and function, triggering a shift from uniform proliferation to a hierarchical organization where the tumor core undergoes a continuous cell turnover.

Overall, this work reveals the existence of self-organization programs that spatially and temporally coordinate cancer cell states. Such self-organization is a hallmark of embryonic tissues (*70, 71*), suggesting that cancer cell dedifferentiation during tumor progression may reactivate developmental programs governing tissue patterning. Based on our findings, we propose that self-organization acts as a fundamental driver of tumor heterogeneity and plasticity, which likely operates synergistically with microenvironmental cues to collectively orchestrate phenotypic transitions during tumor growth, metastasis, therapy resistance and relapse.

## Author contributions

CPG conceived the project, performed most of the experiments, analyzed the data and wrote the manuscript. DBB and EH developed the theoretical model. DBB developed and performed simulations and wrote the manuscript. MdL performed the tumor compression experiments under the supervision of JG-G. SR performed mouse organoid injections and immunostainings. RG fabricated and characterized the polyacrylamide beads under the supervision of SG and JG. MB assisted with single-cell sequencing, FP-F performed immunostainings, NF and RB performed experiments and supported the development of the project. AF, MvdN, MG and JvR contributed technical expertise, materials and discussions. DMV supervised the project and helped writing the manuscript. All authors revised the manuscript.

## Acknowledgements

We acknowledge all the members of the Vignjevic team for support and discussions; the Guillermet-Guibert team and specially Silvia Arcucci and Nicole Therville for assistance with tumor compression experiments and for their hospitality; Morgan Delarue for discussions; Xavier Trepat, Pere Roca-Cusachs and their teams for personal support and invaluable help; Frederic de Sauvage for providing the organoids; the van Rheenen team and specially Maria Azkanak and Dimitrios Laskaris for assistance and for their hospitality; and Jordi Guiu and his team for support and discussions. We thank the platform of Cell and Tissue Imaging (PICT-IBiSA), Institut Curie, member of the national infrastructure France-BioImaging (https://ror.org/01y7vt929) supported by the French National Research Agency (ANR-24-INBS-0005 FBI BIOGEN). We also thank the Institut Curie sequencing platform, especially Sonia Lameiras; Genosplice, especially Noémie Robil and Ariane Jolly for the analysis of single cell sequencing; and the IRB histopathology facility. CPG was funded by an ARC long-term postdoctoral fellowship, GEFLUC, LabEx Cell(n)Scale and the European Union’s Horizon 2020 Marie Skłodowska-Curie grant No 101210382. D.B.B. was supported by the NOMIS foundation as a NOMIS fellow, by the European Molecular Biology Organization (Postdoctoral Fellowship ALTF 343-2022), and by the Austrian Academy of Sciences through an APART-MINT Fellowship. MDL was funded by INCa PLBIO and ITMO MCMP. RG was funded by the European Unions Horizon 2020 research and innovation programs No. 953121 (project FLAMIN-GO). RG, SG, and JG acknowledge core institutional funding from the Max-Planck Society. This work received funding from European Research Council (ERC), under the grant agreement CoG 772487.

