## Supplementary material for "Self-organization of tumor heterogeneity and plasticity"

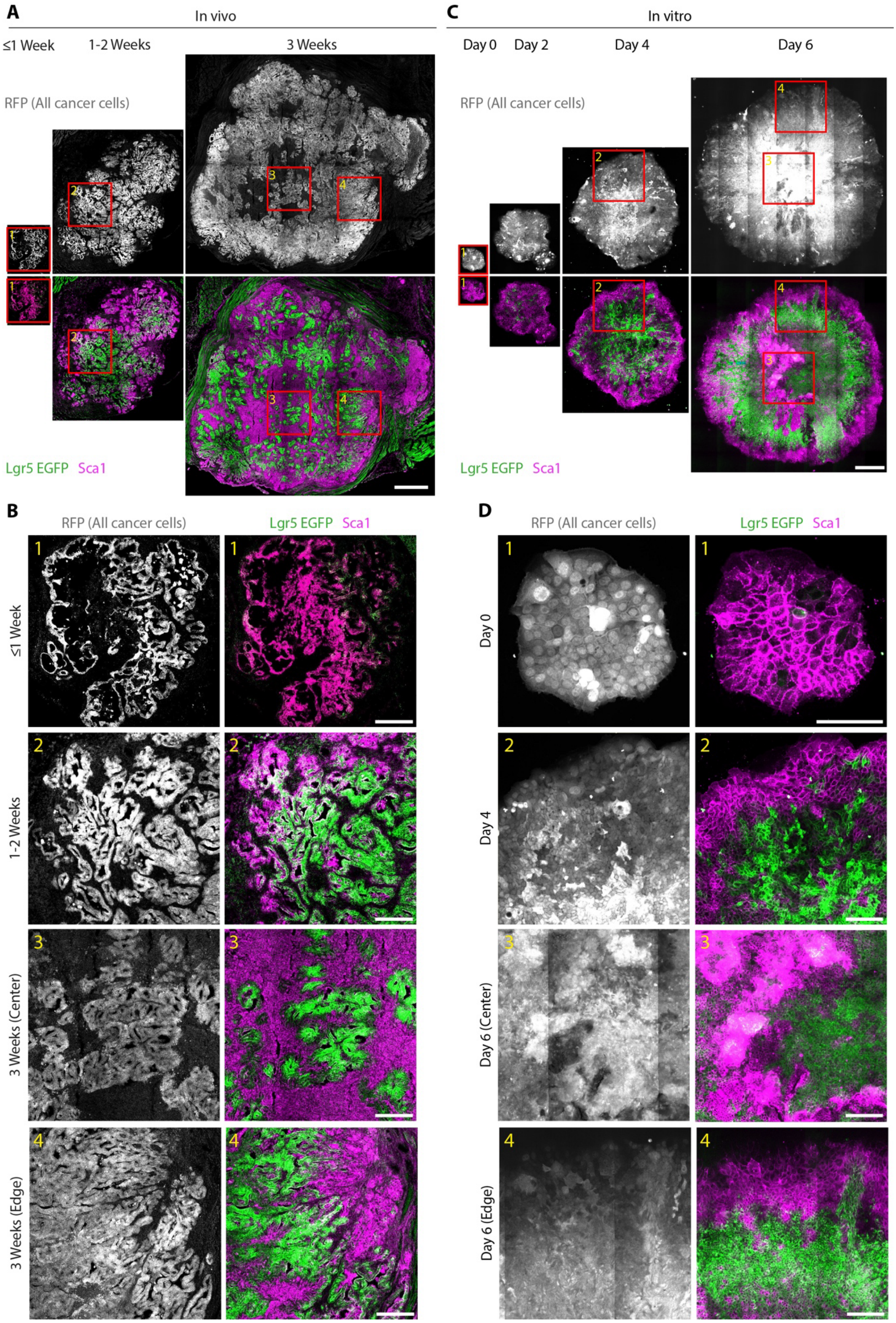


**Fig. S1. High magnification images of subcutaneous tumors and expanding organoids.** **(A-B)** Full tumor section imaging (A) and insets (B) of RFP (all cancer cells) and Lgr5 EGFP-Sca1 at different times after organoid injection. Numbered red squares in A label the insets shown in B. Scalebar = 1000 μm (A) and 250 μm (B, inset). **(C-D)** Full organoid imaging (C) and insets (D) of RFP (all cancer cells) and Lgr5 EGFP-Sca1 at different times after seeding. Numbered red squares in C label the insets shown in D. Scalebar =300 μm (C) and 100 μm (D, inset).

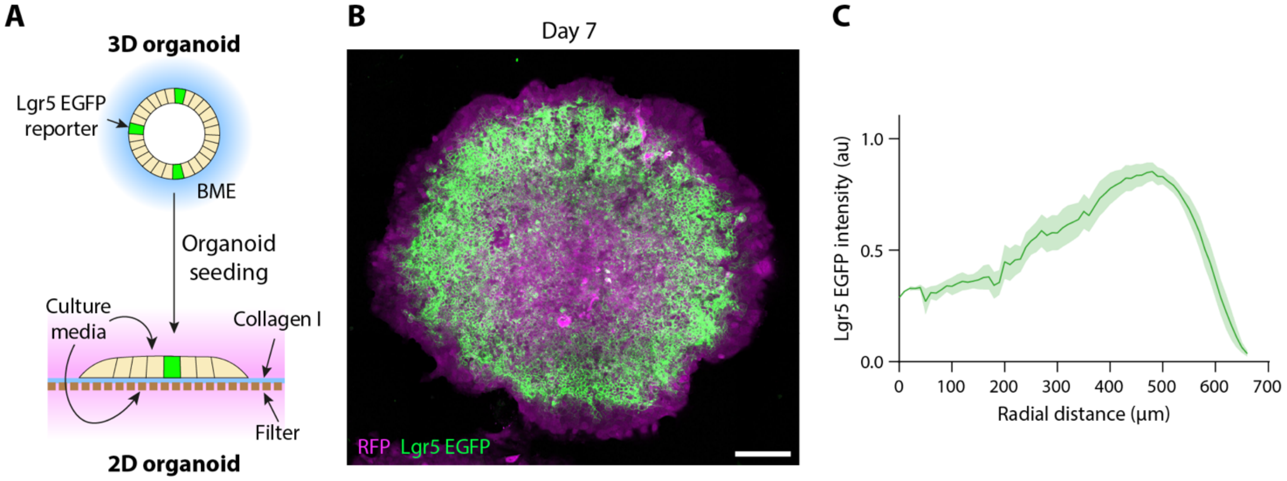


**Fig. S2. Organoid radial patterning on permeable substrates.**
**(A)** Scheme of the experimental setup. **(B)** Representative Lgr5 EGFP and RFP image in an expanding organoid 7 days after seeding on a filter. Scalebar = 100 µm. **(C)** Organoid radial Lgr5 profile at day 7 after seeding on a filter. Mean ± SEM of n=10 organoids from N=2 independent experiments.

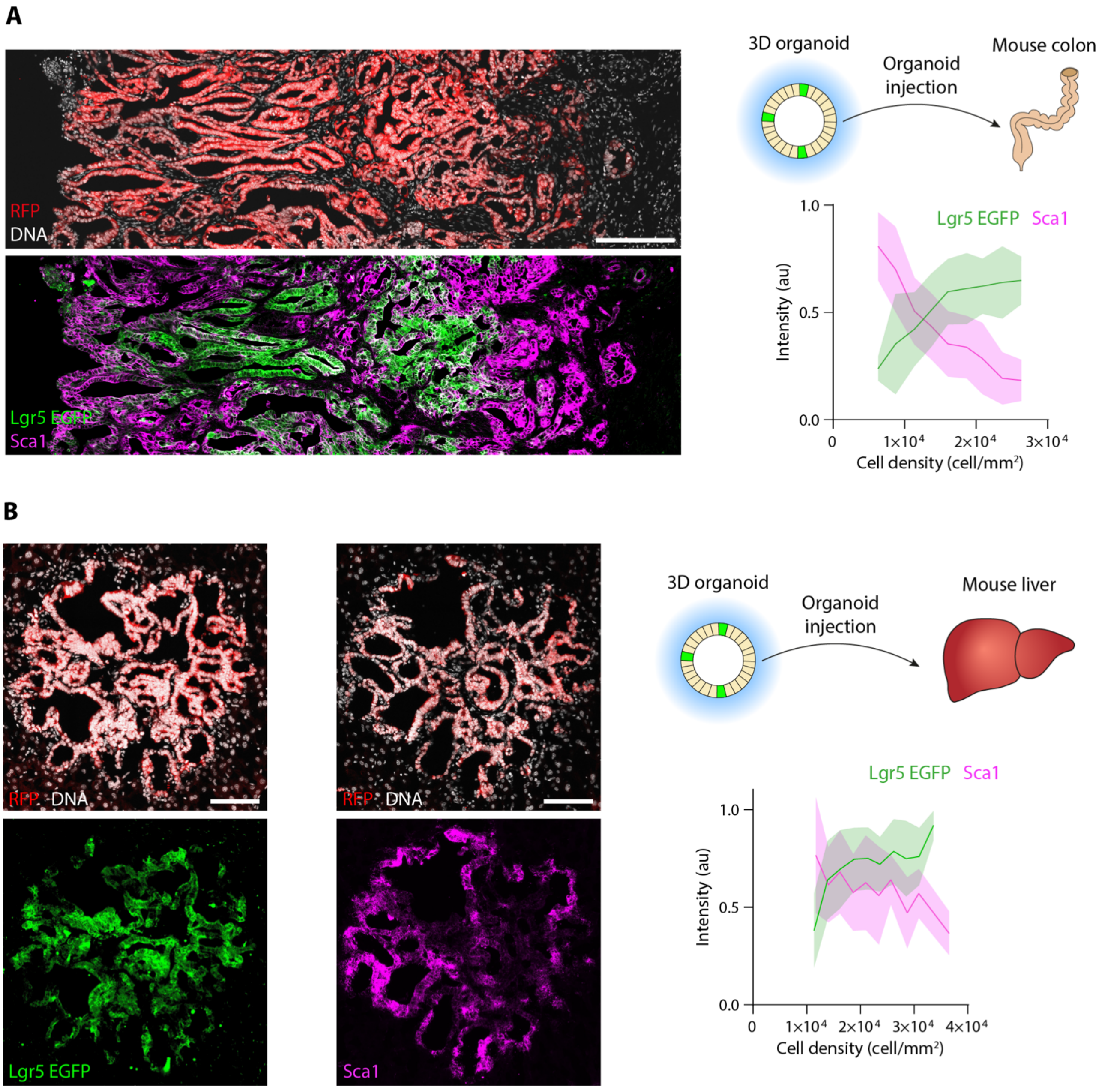


**Fig. S3. State-density relation in colon and liver tumors.**
**(A)** Left: Imaging of RFP (all cancer cells) and DNA (Top), and Lgr5 EGFP and Sca1 (bottom) in an orthotopic colon tumor. Scalebar = 200µm. Right: Scheme of the organoid transplantation (top). Intensity of Lgr5 EGFP and Sca1 as a function of local cell density. Mean ± SD of n=334 local measurements from N=4 tumors. **(B)** Left: Imaging of RFP (all cancer cells) and DNA (Top), and immunostainings of Lgr5 EGFP and Sca1 (bottom) in liver tumor. Note that, due to liver autofluorescence, Sca1 and Lgr5 EGFP (anti-EGFP) had to be stained separately in independent sections. Scalebar = 100µm. Right: Scheme of the organoid transplantation (top). Intensity of Lgr5 EGFP and Sca1 as a function of local cell density. Mean ± SD of n=170 (Lgr5) and 128 (Sca1) local measurements in 12 (Lgr5) and 9 (Sca1) tumors from N=2 mice.

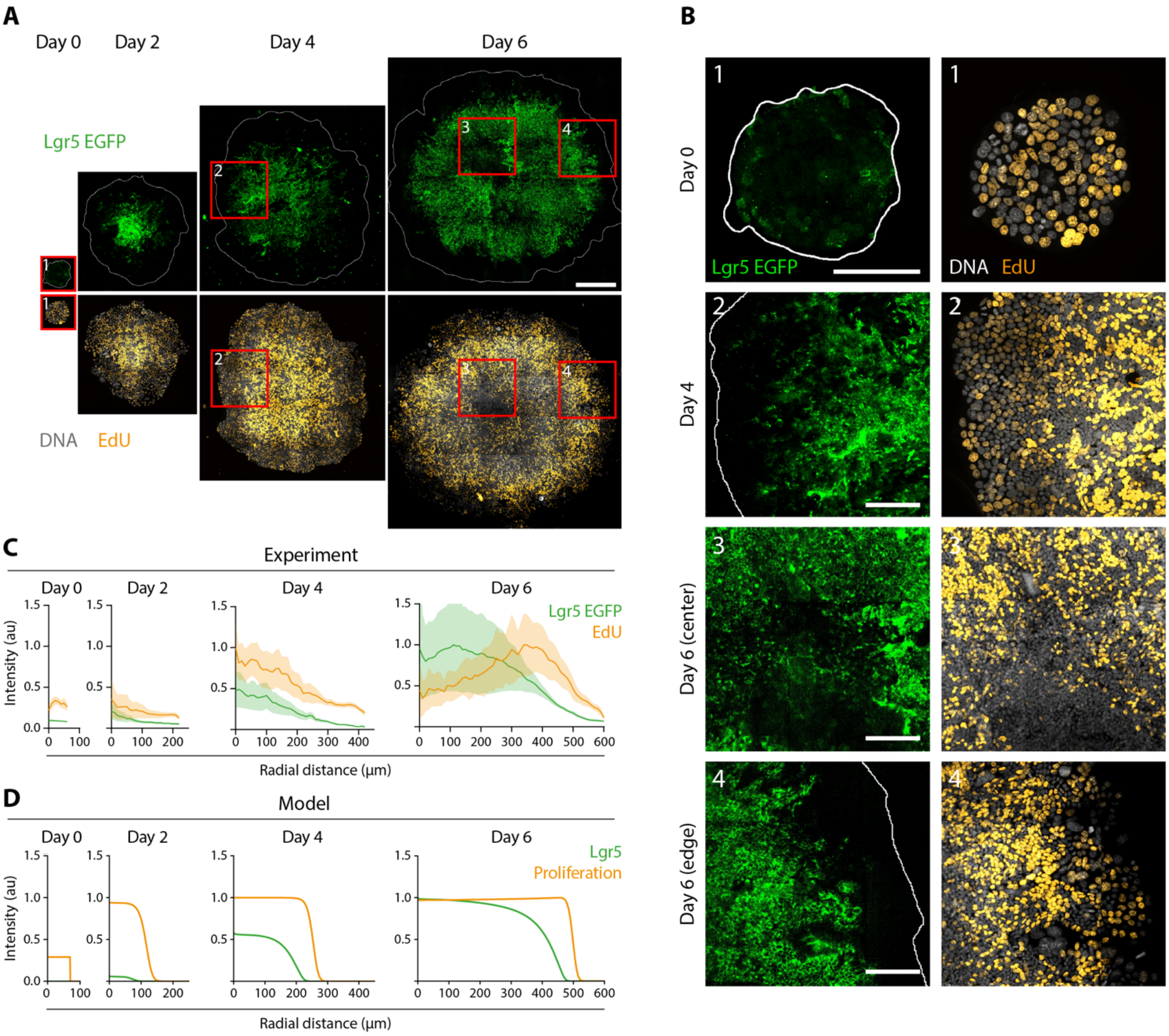


**Fig. S4. Evolution of cell proliferation in expanding organoids.**
**(A-B)** Full organoid imaging (A) and insets (B) of Lgr5 EGFP, EdU and DNA at different times after seeding. Color squares in A label the insets shown in B. Scalebar =300 µm (A) and 100 µm (B, inset). **(C-D)** Time evolution of radial profiles of Lgr5 EGFP and proliferation (EdU) in experiments (C) and model simulations (D). Mean ± SEM of n=8 (day 0), 8 (day 2), 7 (day 4), 3 (day 6) organoids from N=2 independent experiments.

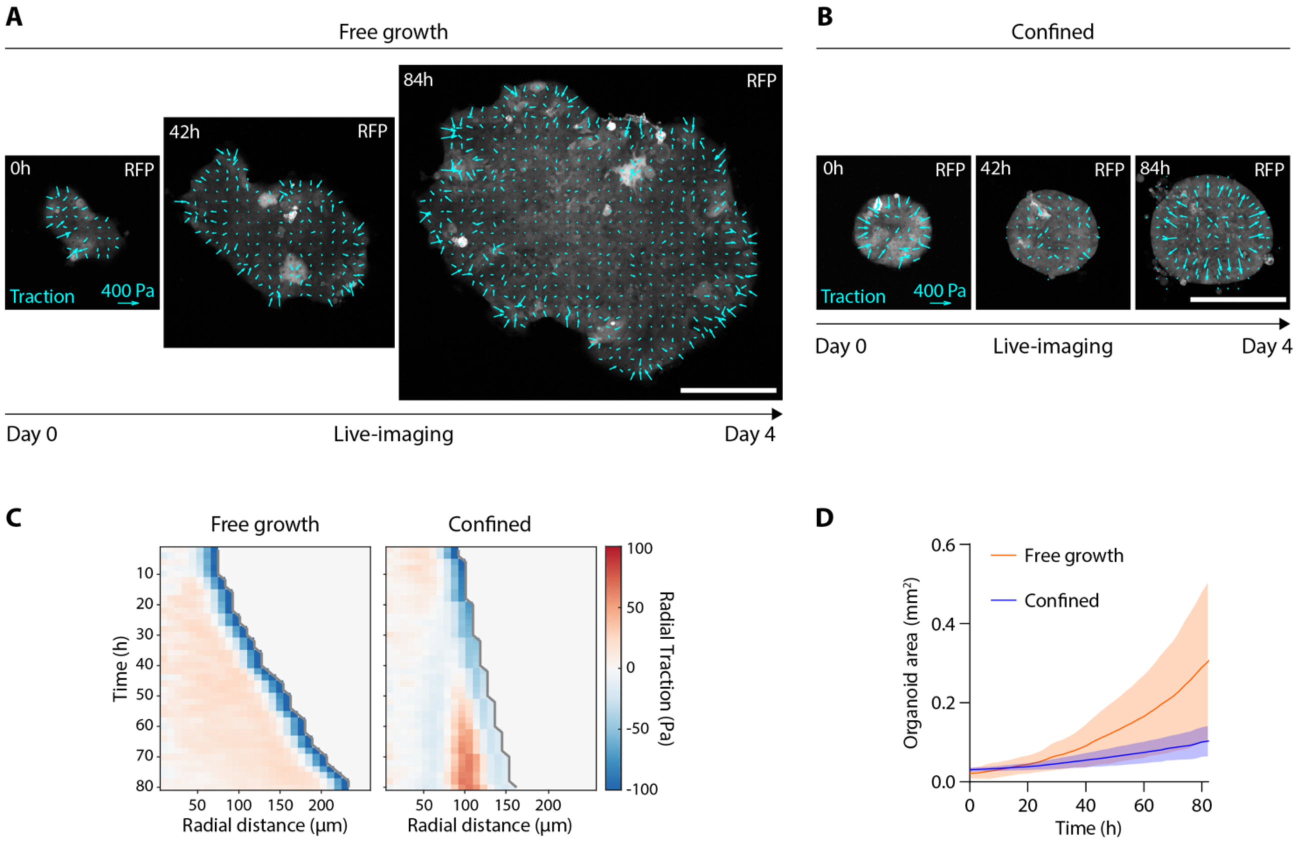


**Fig. S5. Traction forces in freely growing and confined organoids.**
**(A-B)** Evolution of traction force maps in freely growing (A) and confined (B) organoids. Scalebar = 200µm. **(C)** Average kymographs of radial tractions in freely growing (left) and confined (right) organoids. Positive tractions point towards the right. Organoids are the same as shown in Fig. 2A-D. n=10 (free growth) and 22 (confined) organoids from N=3 (free growth) and 4 (confined) independent experiments. **(D)** Evolution of monolayer area in freely growing and confined organoids. Mean ± SD.

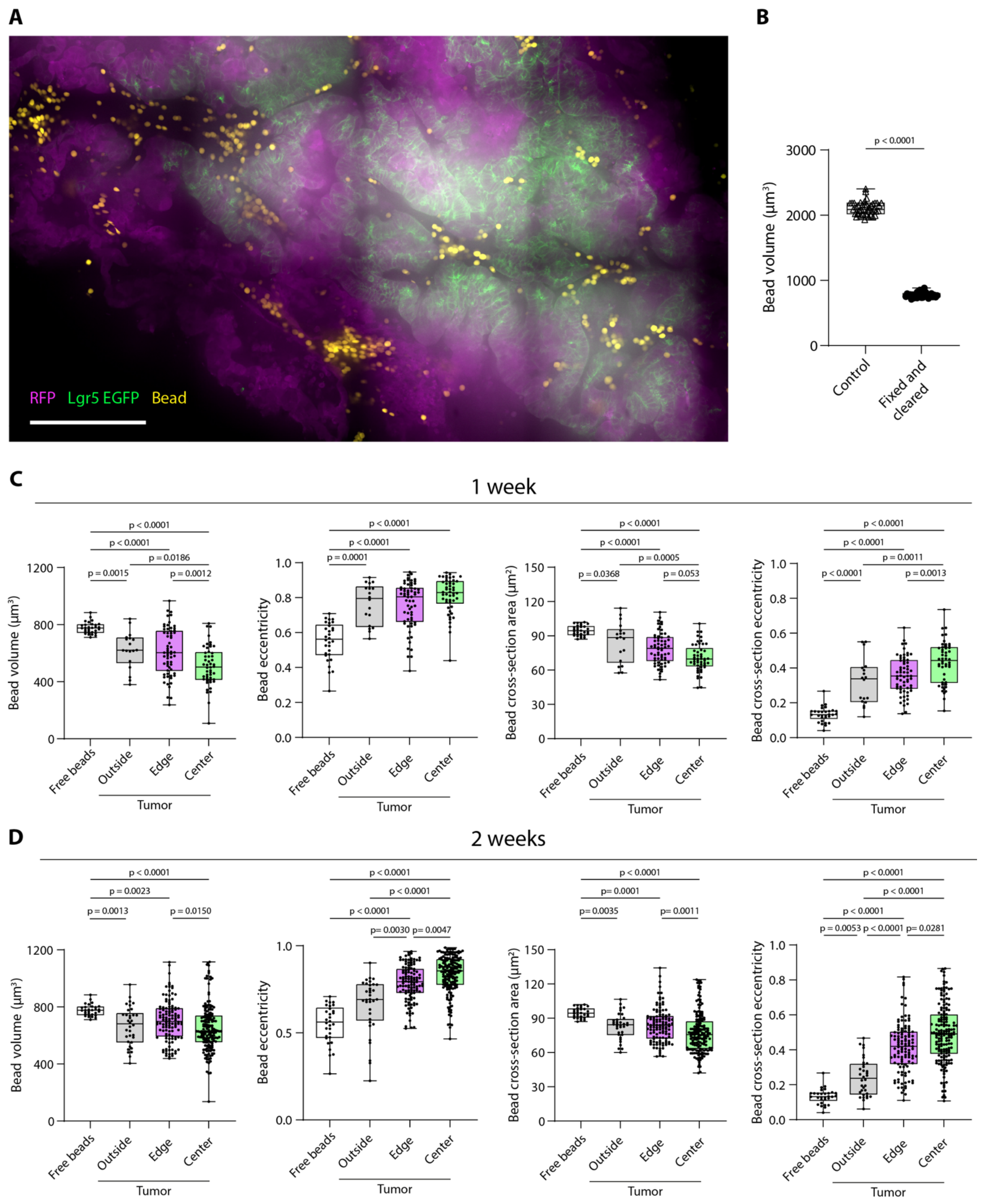


**Fig. S6. Tumor stress measurements using polyacrylamide beads as force sensors.**
**(A)** Representative image of a tumor expressing RFP (all cancer cells) and EGFP (Lgr5+ cells) containing polyacrylamyde beads integrated. **(B)** Bead volume quantification in free beads before (Control) and after fixation and clearing (fixed and cleared). Unpaired t-test. **(C-D)** Bead volume, eccentricity, cross-sectional area and cross-sectional eccentricity at different tumor locations in tumors harvested for 1 (C) and 2 (D) weeks. Normality test (Kolmogorov-Smirnov) was first perform. If all conditions followed a normal distribution (1 week: volume, cross-section area, cross-section eccentricity) one-way ANOVA test was performed. For the other graphs, Kruskal-Wallis test was performed.

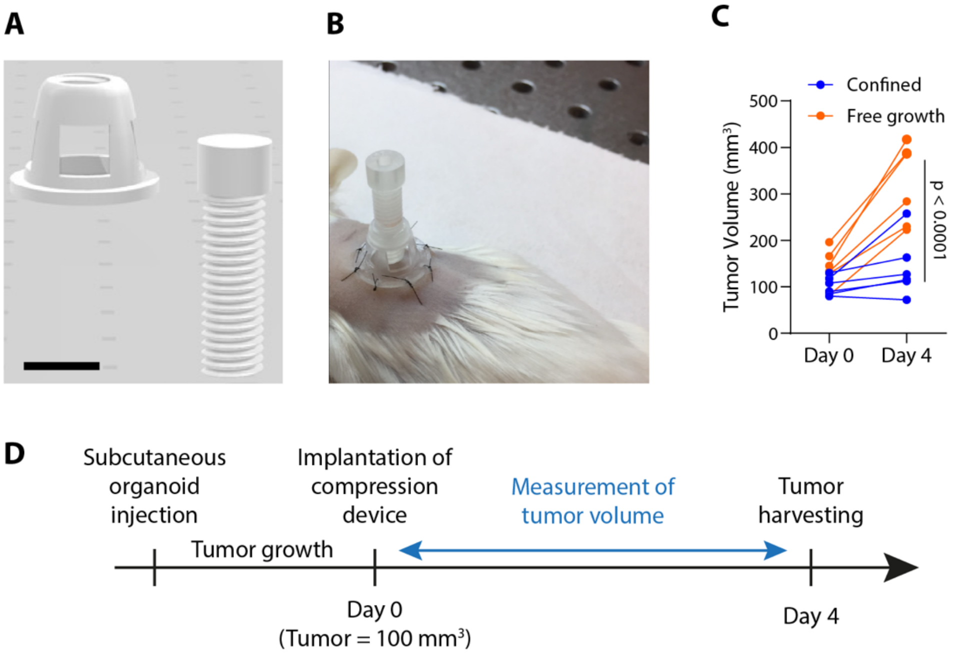


**Fig. S7. Compression device slows down tumor growth**
**(A)** Image of the compression device showing the holder (left) and the screw (right). Scalebar = 5 mm. **(B)** Installation of the compression device on mouse subcutaneous tumors. **(C)** Tumor volume before and after 4 days of compression. N= 5 (free growth) and 6 (confined) tumors. All points are shown. Two-way ANOVA with repeated measures and multiple comparisons. **(D)** Experimental design to confine tumor growth for 4 days. Compression starts (Day 0) when the tumor reaches 100 $mm^3$.

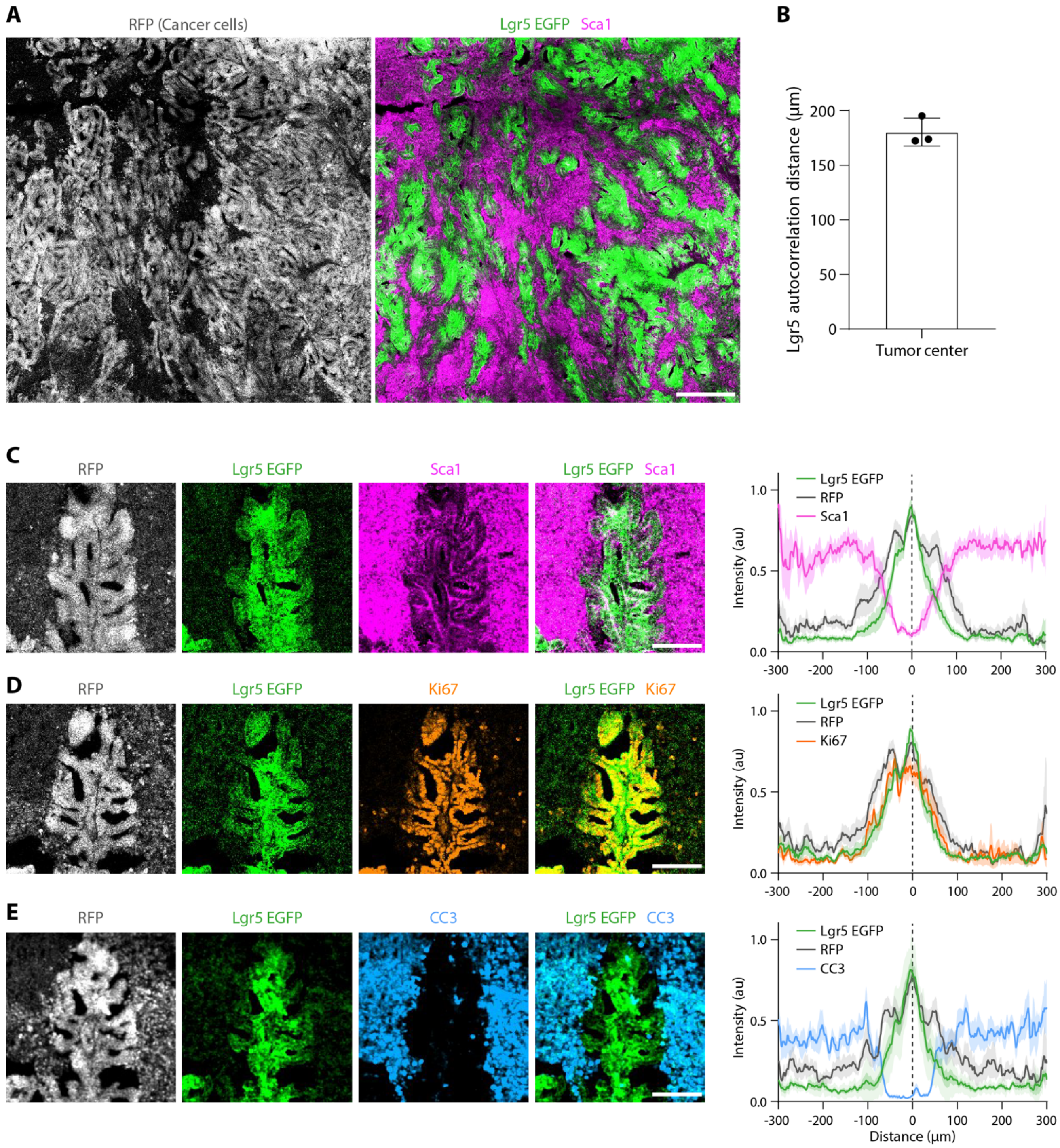


**Fig. S8. Periodic hierarchical units with a characteristic width at the tumor apoptotic core.** **(A)** Representative image of the core of a subcutaneous tumor expressing RFP (left) and Lgr5 EGFP (right) stained for Sca1. Scalebar = 300 µm. **(B)** Average distance between Lgr5 EGFP autocorrelation peaks at the tumor core. Points represent independent tumors (N=3), bar represents Mean ± SD. **(C-E)** Left: representative image of an Lgr5 stripe at the tumor core stained for Sca1 (C), Ki67 (D) and cleaved-caspase 3 (CC3, E). Images show the same strip in consecutive cuts. Right: average linescan of Lgr5 EGFP, RFP, Sca1 (C), Ki67 (D) and CC3 (E) intensity as a function of the distance to the Lgr5 stripe center (distance =0). Mean± SD of n=9 stripes from N=3 tumors. Scalebar = 100 µm.

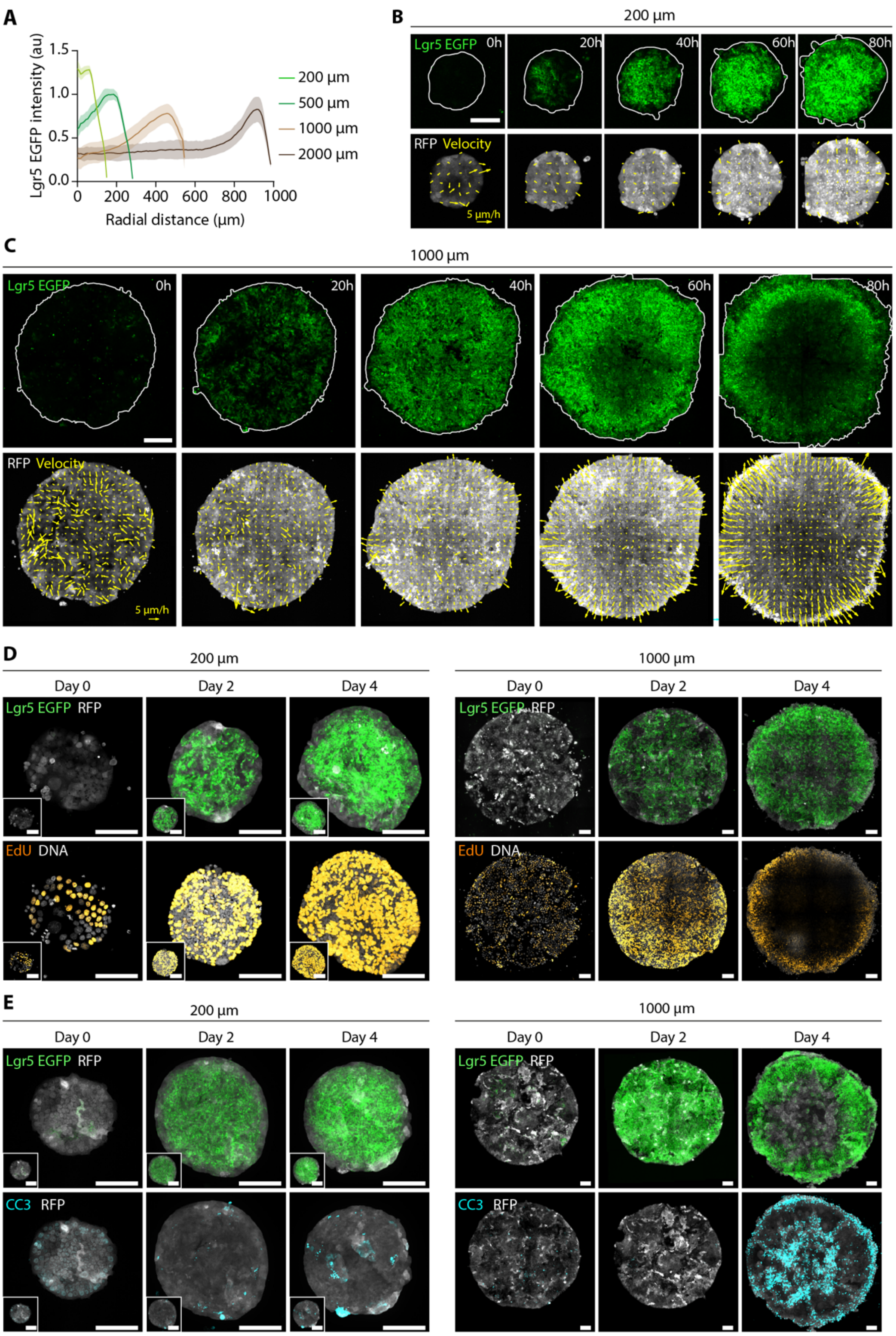


**Fig. S9. Supplementary data for figure 3.**
**(A)** Not-normalized radial Lgr5 EGFP intensity profile in organoids patterned to different sizes at day 4. Note that the Lgr5 EGFP profile in Fig. 3A is normalized to better visualize the constant width of the Lgr5 compartment. **(B-C)** Live imaging of Lgr5 EGFP (top) and RFP with overlayed velocity maps (bottom) for small (B) and large (C) organoids. Scalebar = 100 µm (B) and 200 µm (C). **(D)** Small (left) and large (right) Lgr5 EGFP and RFP expressing organodis (top) stained for EdU and DNA (bottom) at different times after seeding. Scalebar = 100 µm (Small organoids are scaled as large organoids in the bottom left white squares). Organoids are the same shown in fig. 3C-D. **(E)** Small (left) and large (right) Lgr5 EGFP and RFP expressing organodis (top) stained for cleaved-caspase 3 (CC3) (bottom) at different times after seeding. Scalebar = 100 µm (Small organoids are scaled as large organoids in the bottom left white squares). Organoids are the same shown in fig. 3C-D.

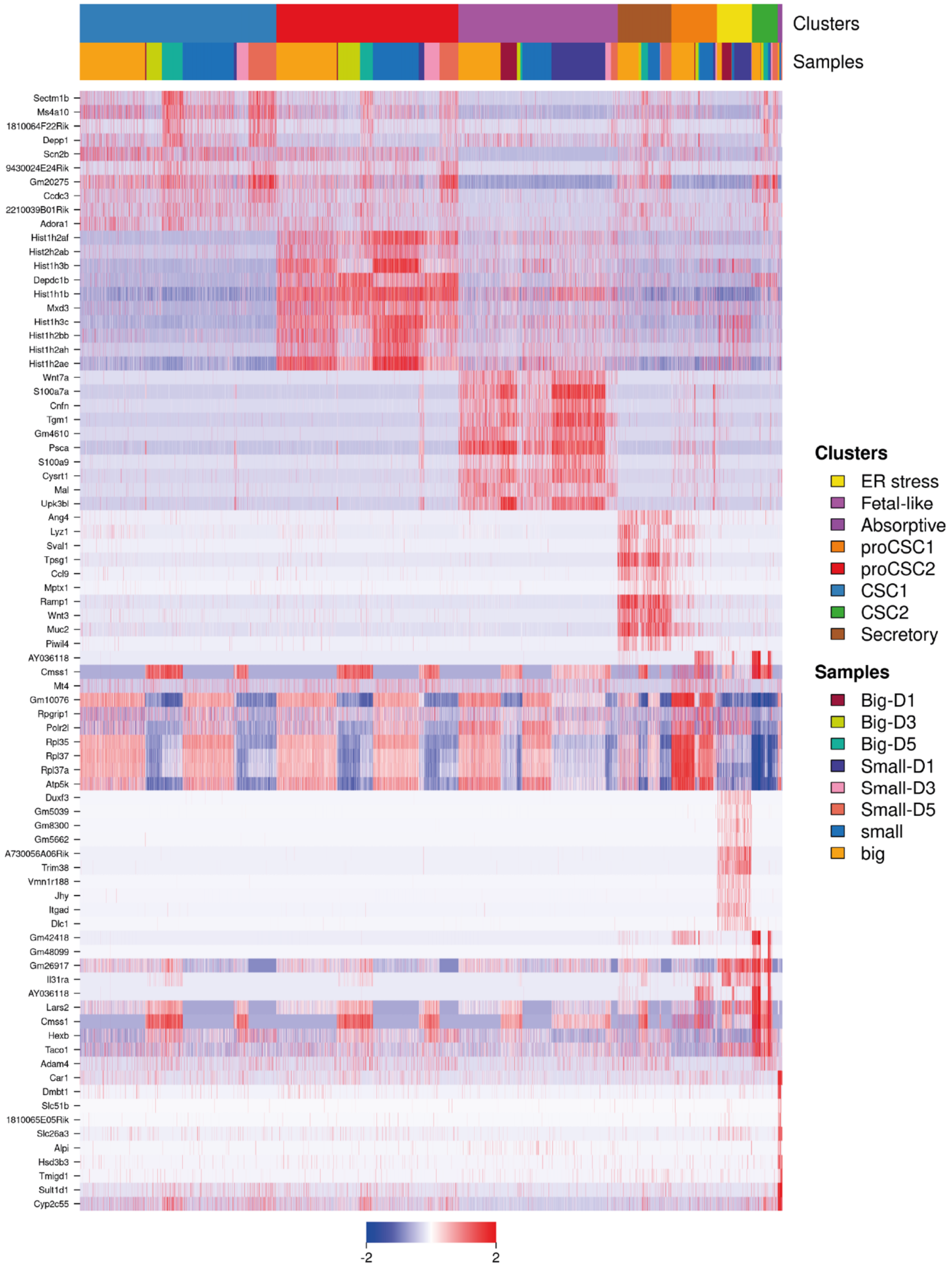


**Fig. S10. Clustering of single cell sequencing data.**
Heatmap of the top 10 marker genes in each cluster.

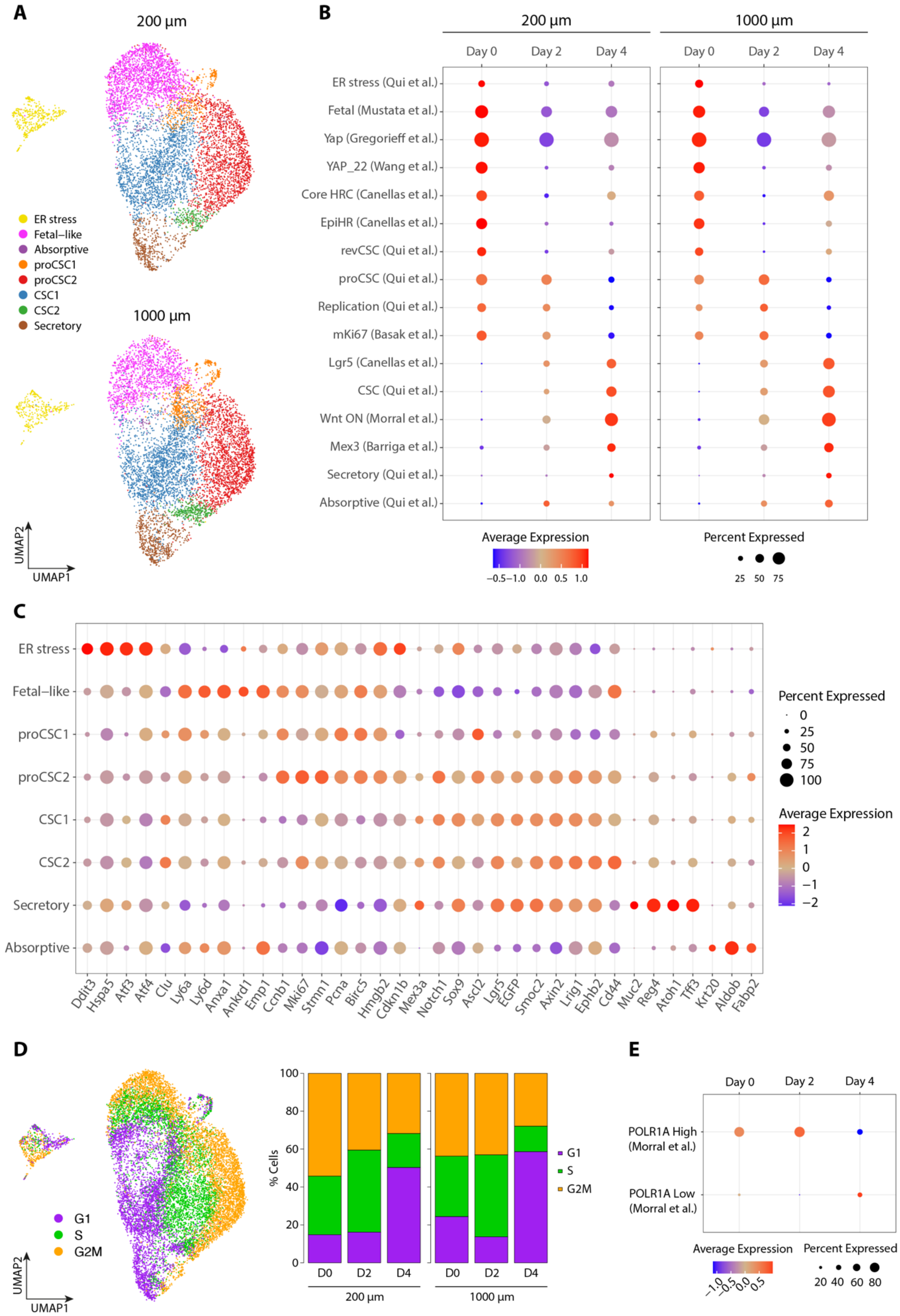


**Fig. S11. Additional single cell sequencing analysis**

**(A)** UMAP showing the time (day 0, 2 or 4) of every single cell of small (200 µm) and large (1000 µm) organoids. **(B)** Score of different signatures at day 0, 2 and 4 in small (left) and large (right) organoids. **(C)** Expression of single genes in each cluster. **(D)** Left: UMAP showing the predicted phase of the cell cycle in each cell. Right: proportion of cells in each phase of the cell cycle per day. **(E)** Score of high (POLR1A high) and low (POLR1A low) biosynthesis signatures at day 0, 2 and 4.

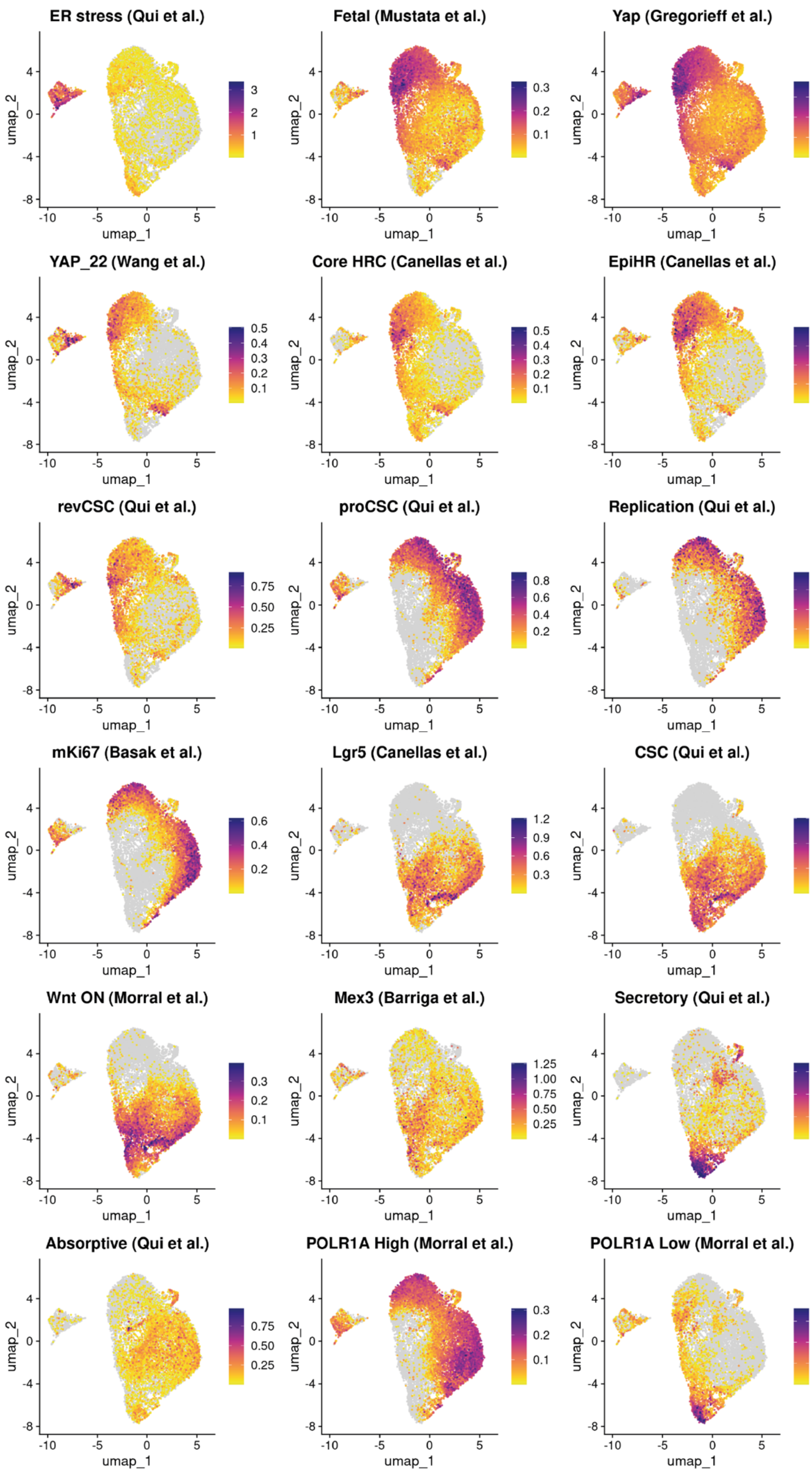


**Fig. S12. Signature score at the single cell level.**
UMAPs showing the score of all signatures used in the study in each single cell.

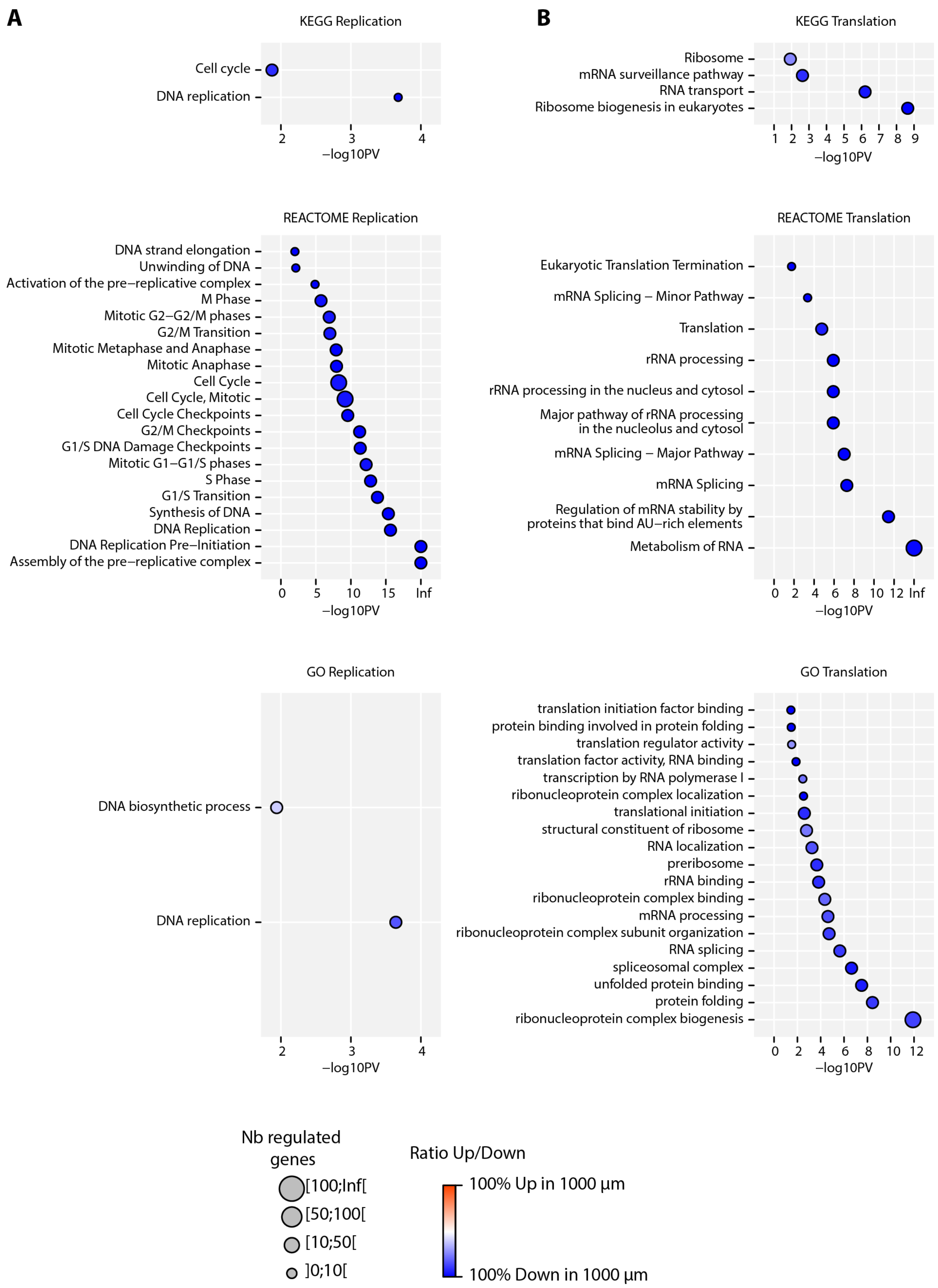


**Fig. S13. Differential expression analysis between small and large organoids at day 4.**
**(A-B)** All significant terms related to replication (A) and RNA metabolism and translation (B) from KEGG (top), REACTOME (middle) and GO (bottom) databases.

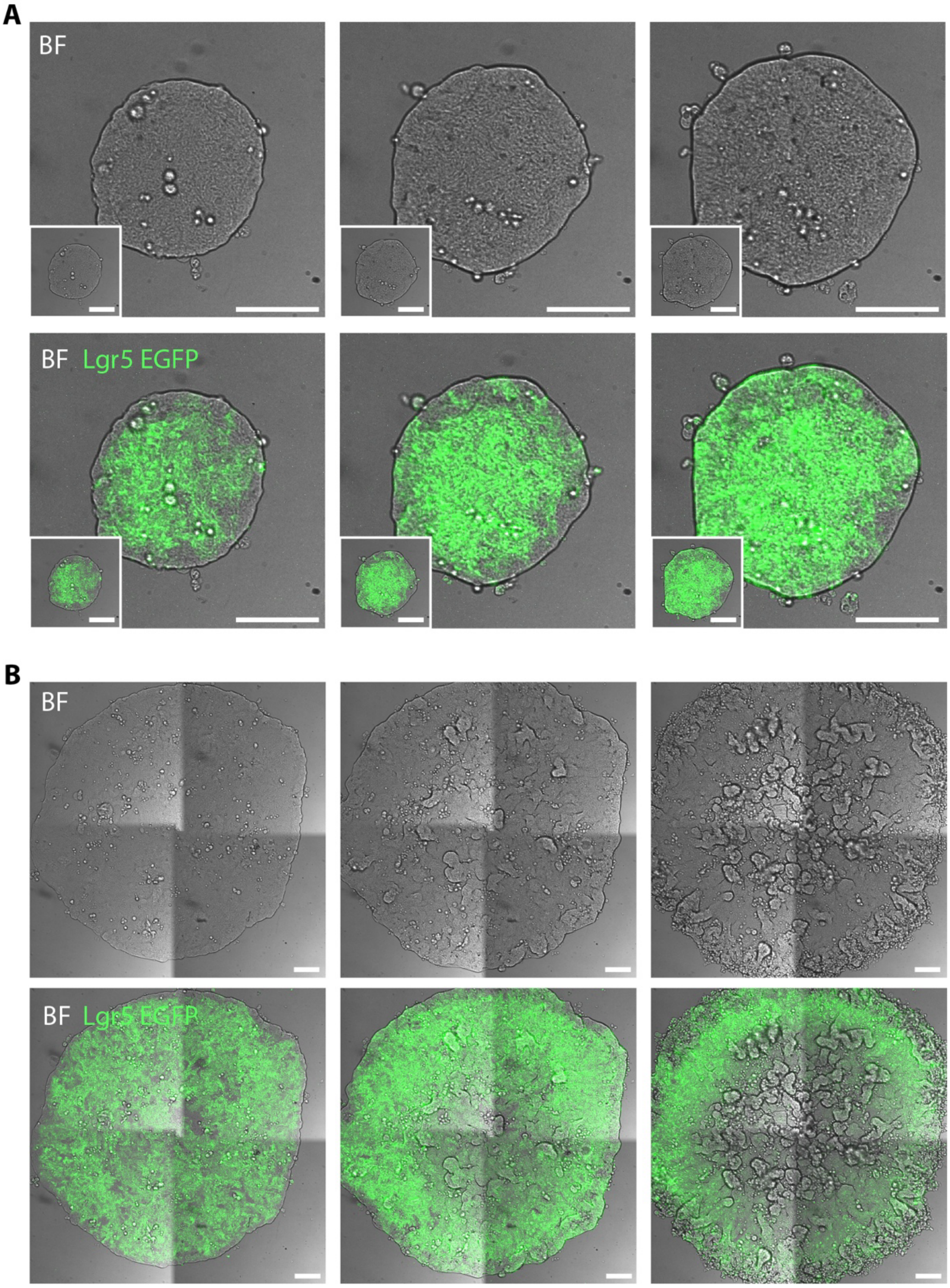


**Fig. S14. Tissue folding upon loss of Lgr5 at the center of large organoids.**
**(A-B)** Brightfield and overlaid Lgr5 EGFP signal from small (200 µm, A) and large (1000 µm, B) organoids. Scalebar = 100 µm (Small organoids are scaled as large organoids in the bottom left white squares).

**Movie captions**

**Movie S1. Organoid spreading velocity.**
Timelapse of organoid spreading (RFP expressed by all cells) with overlaid velocity maps (yellow vectors). Scalebar = 100 µm. Vector scale (bottom right) = 10 µm/h.

**Movie S2. Coevolution of Lgr5 EGFP and intercellular stresses in spreading and confined organoids.**
Top: Lgr5 EGFP time evolution in freely growing (left) and confined (right) organoids. White lines label the organoid contour. Bottom: intercellular stress maps overlaid on organoid fluorescent images (RFP) in freely growing (left) and confined (right) organoids. Scalebar = 200 µm.

**Movie S3. Time evolution of traction forces in freely growing and confined organoids.**
Traction forces (cyan vectors) overlaid on organoid fluorescent images (RFP) for freely growing (left) and confined (right) organoids. Scalebar = 200 µm. Vector scale (bottom right) = 400Pa.

**Movie S4. Lgr5 dynamics in small (200 µm) and large (1000 µm) organoids.**
Lgr5 EGFP (left) and merged Lgr5 and RFP images (right) for small (top) and large (bottom) organoids. Scalebar = 200 µm.

**Movie S5. Velocity of small (200 µm) and large (1000 µm) organoids.**
Velocity maps (yellow vectors) overlaid on small (left) and large (right) organoid images (RFP). Scalebar = 200 µm. Vector scale = 5 µm/h.

**Movie S6. RFP dynamics in small (200 µm) and large (1000 µm) organoids.**
RFP (left) and merged Lgr5 and RFP images (right) for small (top) and large (bottom) organoids. Scalebar = 200 µm.

**Movie S7. Lgr5 EGFP decrease at the center of large organoids is paralleled by tissue folding.**
Bright-field (left) and merged Lgr5 and bright-field images (right) for small (top) and large (bottom) organoids. Scalebar = 200 µm.

**Theory note: Development of a physical model of cancer stem cell organoid patterning**

Here we present a theoretical model of the coupled dynamics of cell proliferation, stress, velocity and Lgr5 dynamics in expanding and confined cancer stem cell organoids. First, we develop a continuum theory that considers the balance of active forces within organoids at the multicellular scale to predict growth and spreading dynamics. Second, we incorporate cell state dynamics into the model using a density-sensitive Lgr5 activation mechanism to predict spatio-temporal patterning dynamics.

**1. Setup of the active fluid model**

We first derive the main equations governing the system, starting from the appropriate conservations laws and constitutive relations. We consider the radial coordinate *r*, and seek to derive the equations governing the cell density profile $\rho(r)$ and the profile of radial velocities $v_r(r)$.

The cell density obeys the conservation law

$$\frac{\partial \rho}{\partial t} = -\frac{1}{r}\frac{\partial (rJ)}{\partial r} + f(\rho) \qquad (1)$$

where the first term is the divergence of a radial flux of cells $J(r)$ and $f(\rho)$ describes the dynamics of cell division and death. The constitutive equation for the flux is given by

$$J = -D\frac{\partial \rho}{\partial r} + \rho v_r \qquad (2)$$

where the first term describes a diffusive flux (random movement of cells with diffusion coefficient $D$), and the second term describes advection of cells by the velocity field.

The velocity obeys conservation of momentum (i.e. force balance), which in the low Reynolds number regime is given by

$$\nabla \cdot \boldsymbol{\sigma} = \zeta \boldsymbol{v} - \boldsymbol{f}_A = 0 \qquad (3)$$

where $\zeta$ is the friction coefficient between the cells and the substrate, $\boldsymbol{\sigma}$ is the stress tensor within the tissue and $\boldsymbol{f}_A$ is an active force generated by cells that acts on the substrate. Based on the observation that there is no significant swirling motion of cells in the organoids, we are solving a radially symmetric problem, we neglect shear stress components ($\sigma_{\theta r} = 0$) and focus on the *r*-component of the force-balance equation, given by

$$\frac{\partial \sigma_{rr}}{\partial r} + \frac{1}{r}(\sigma_{rr} - \sigma_{\theta\theta}) = \zeta v_r - f_A(r) \qquad (4)$$

where $\sigma_{rr}$ is the radial stress and $\sigma_{\theta\theta}$ is the hoop stress.

At long-time scales compared to the dynamics of cell-cell rearrangements, which is valid here as we study dynamics of spreading over days, tissues can be described as active viscous media[1–4]. With radially symmetric flow, the constitutive equations for the stress are then given by

$$\sigma_{rr} = \eta\frac{\partial v_r}{\partial r} - P \qquad (5)$$

$$\sigma_{\theta\theta} = \eta\frac{v_r}{r} - P \qquad (6)$$

where the first term in (5) represents the radial viscous stress within the cell layer with viscosity $\eta$, and the first term in (6) represents the hoop strain rate from circumferential expansion of the tissue. $P$ is the pressure, which we expand linearly in cell density[1]:

$$P = \chi(\rho - \rho_{eq}) \qquad (7)$$

where $\chi$ is a compressibility parameter and $\rho_{eq}$ is the equilibrium cell density. Note that $\rho_{eq}$ is irrelevant for the force balance equation, and thus for the governing equation of velocities, as it disappears when taking gradients of pressure, yet it is important to set the scale of tension vs compressive stresses when evaluating the radial stress. Furthermore, for simplicity, we absorbed any additional stress terms due to compressible volumetric deformation (determined by the bulk viscosity and the divergence of the velocity field) into the effective pressure term.

Equations (1)-(5) describe the time evolution of the cell density and the corresponding velocity field, and can be combined into the two governing partial differential equations

$$\frac{\partial \rho}{\partial t} = D\frac{1}{r}\frac{\partial}{\partial r}\left(r\frac{\partial \rho}{\partial r}\right) - \frac{1}{r}\frac{\partial (r\rho v_r)}{\partial r} + f(\rho) \tag{8}$$

$$0 = \eta\frac{\partial^2 v_r}{\partial r^2} + \frac{\eta}{r}\frac{\partial v_r}{\partial r} - v_r\left(\zeta + \frac{\eta}{r^2}\right) - \chi\frac{\partial \rho}{\partial r} + f_A(r) \tag{9}$$

To fully specify the model, we have to choose the form of the active force and the cell proliferation term. For the active force, we find that a constant pulling force at the edge simply rescales the global tension and is therefore absorbed into the pressure term[5]; for simplicity we therefore take $f_A(r) = 0$. For the proliferation dynamics, we assume logistic growth with a growth rate $k_d$ and a critical density $\rho^*$

$$f(\rho) = k_d \rho(1 - \rho/\rho^*) \tag{10}$$

We consider no flux boundary conditions for the cell density at $r = 0$ and $r = R_\infty$:

$$\frac{\partial \rho(r = 0, t)}{\partial r} = \frac{\partial \rho(r = R_\infty, t)}{\partial r} = 0 \tag{11}$$

and clamped boundary conditions for the radial velocity at $r = 0$ and $r = R_\infty$:

$$v_r(r = 0, t) = v_r(r = R_\infty, t) = 0 \tag{12}$$

For the initial conditions, we use

$$\rho(r, t = 0) = \rho_0 \qquad v_r(r, t = 0) = 0 \tag{13}$$

Where the density initial condition is set to zero outside the tissue. Since the confined systems also exhibit a non-negligible outward expansion, which is central for setting the length-scale of stress decay into the tissue, we use these boundary conditions for both free and confined systems, rather than setting closed boundary conditions for confined systems. Instead, we implement free vs confined dynamics by adapting the friction coefficient $\zeta$, i.e. $\zeta = \zeta_{\text{confined}}$ and $\zeta = \zeta_{\text{free}}$.

### 2. Parameter fitting

*2.1. Density dynamics*

We first fit the parameters of cell proliferation using the evolution of cell density in free and confined systems. By plotting normalized growth rate against normalized cell density, we can directly fit the logistic growth equation (Eq. (10)) and infer the best fit parameters in units of the initial cell density $\rho_0$ : $k_d = 1\ \rho_0/\text{day}$ and $\rho^* = 12.7\rho_0$. For simplicity, we further assume that random cell motility within the tissue is negligible and thus take $D = 0.01$. This fully parameterizes Eq. (10).

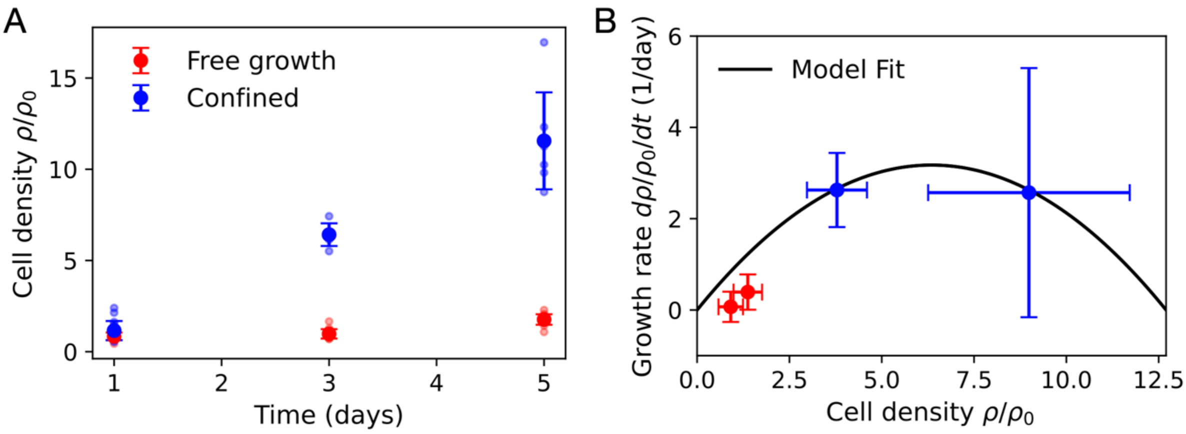


**Fig. M1: Fitting cell density growth. A**, Dynamics of normalized cell density in free (red) and confined (blue) systems. **B**, Normalized growth rate (measured as discrete derivative from data in A) vs normalized cell density. Black line represents the model fit of Eq. (10) with best fit parameters $k_d = 1\ \rho_0/\text{day}$ and $\rho^* = 12.7\rho_0$.

*2.2. Velocity dynamics*

To fully parameterize the active fluid model, we next determine the parameters of Eq. (9): $\eta, \chi, \zeta_{\text{free}}, \zeta_{\text{confined}}$. For this, we fit the dynamics of edge expansion of the free growth system to determine $\eta, \chi, \zeta_{\text{free}}$ by fitting the expansion of the colony edge over time (Fig. M2A), as well as the spatial and temporal dependence of radial velocities (Fig. M2B-C). This results in best-fit parameters $\eta = 0.7, \chi = 0.1, \zeta_{\text{free}} = 0.05$.

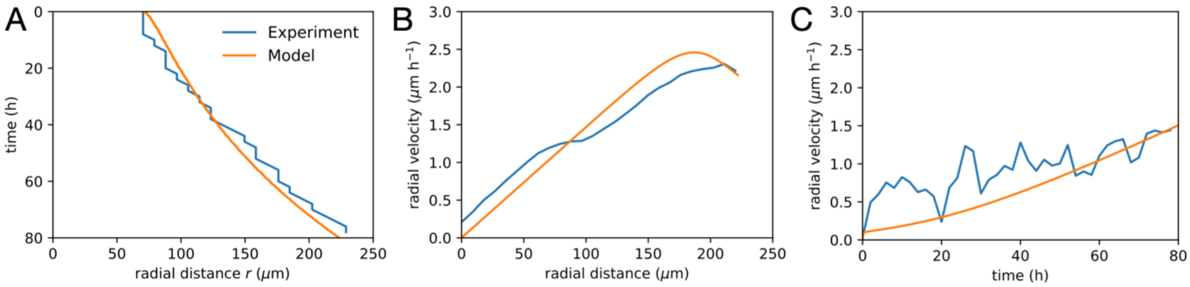


**Fig. M2: Fitting tissue expansion dynamics of free growth systems. A,** Location of the colony edge over time for experiment (blue) and model (orange). In the model, the edge is defined as the location at which the cell density falls below 5% of the initial cell density, i.e. $\rho < 0.05\rho_0$. **B**, Radial velocity at $t$ = 80h as a function of radial distance. **C**, Radial velocity averaged over the whole system as a function of time.

*2.3. Prediction of intercellular stress dynamics in expanding vs confined systems*

Having constrained the key parameters of the active fluid model, we make predictions for how the dynamics are expected to change in the presence of confinement. To incorporate the slow expansion of confined systems in the model, we use an effective implementation of confinement by increasing the friction coefficient, i.e. $\zeta_{\text{confined}} \gg \zeta_{\text{free}}$. While simulations with complete confinement, i.e. no-flux boundary conditions at the confinement radius $r = R$, give qualitatively similar results for the transition to compressive stress, the slow expansion gives rise to a boundary layer of higher stress and lower density, as observed experimentally. Our model then predicts that the intercellular stress becomes compressive in confined systems while it is still tensile in free growth systems, due to higher cell density in confined systems (Fig. M3A-D).

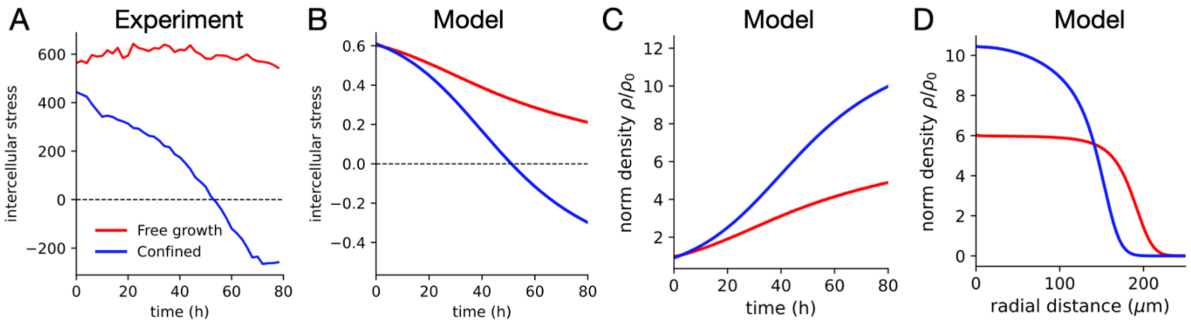


**Fig. M3: Fitting intercellular stress dynamics. A,** intercellular stress in free growth (red) and confined (blue) systems as a function of time. **B**, equivalent plot for the model. For both model and experiment, the stress is averaged over the whole colony, excluding the expanding rim in confined colonies to highlight the compressive nature of the stress in the center. Parameters determined from Figs. M1, M2 are used in addition to equilibrium cell density that controls the switch between extensile and compressive stress, $\rho_{eq} = 7$.

#### 3. Density-dependent cell state model

To determine whether density-dependent cellular states of Lgr5+ vs Sca1+ are sufficient to explain the experimentally observed time-dependent patterning of Lgr5 and Sca1, we first consider a density-dependent cell state model in which Lgr5 is positively regulated by cell density:

$$\frac{dL}{dt} = A(\rho; K_L, h) - d_L L \tag{14}$$

where $A(x; K, h)$ is an activating Hill function

$$A(x; K, h) = \frac{x^h}{x^h + K^h} \tag{15}$$

The Lgr5 dynamics are simulated with initial conditions $L(x, t = 0) = 0$.

To predict Sca1-dynamics, we hypothesize that Sca1+ is a default cell state that is assumed whenever cells are Lgr5- (Fig. M4). As a minimal model for this, we assume the Sca1 production term to be given by

$$\frac{dS}{dt} = 1 - A(\rho; K_L, h) - d_S S \tag{16}$$

where we use initial condition $S(x, t = 0) = 1$ since cells are initially Sca1+.

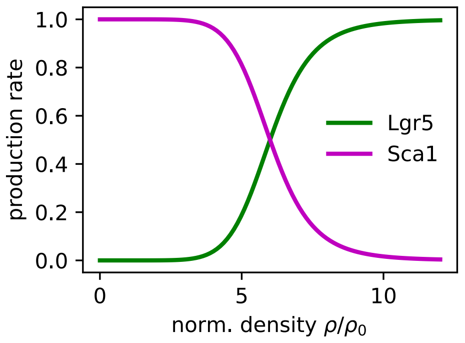


**Fig. M4: Lgr5 and Sca1 production terms assumed in the density-dependent cell state model.**

The parameters of the density-dependent cell state model are fit to the experimentally measured cell state dynamics in free growth and confined systems to account for the timing of the onset of Lgr5 production. This model correctly captures the slower and weaker Lgr5 production in free growth systems compared to confined systems, and the accompanying loss of Sca1 in the center of free and confined systems (Fig. M5A, B). Note that throughout the paper, for all intensity-based measures (Lgr5-GFP, Sca1, RFP, EdU), the predicted intensities are multiplied by the local tissue density, to account for the fact that denser regions appear brighter in fluorescence images. Importantly, it also predicts a decrease the width of the Sca1 rim at the tissue edge in confined systems (Fig. M5B), which is due to the density gradient being less steep than in confined systems.

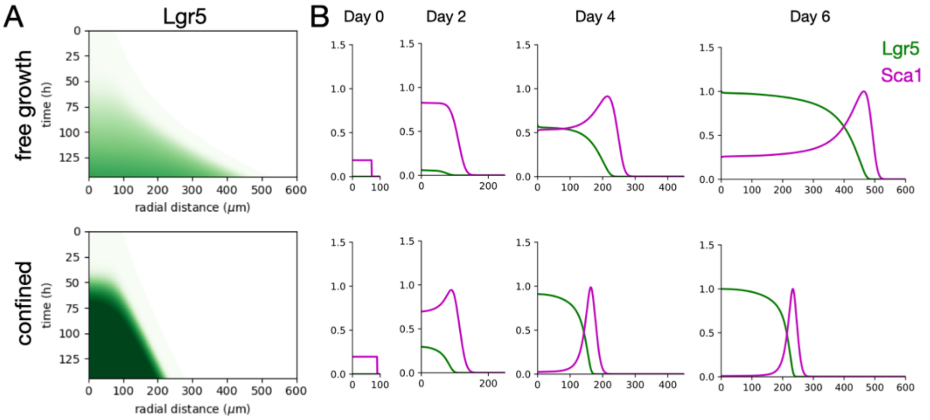


**Fig. M5: Lgr5 and Sca1 dynamics predicted by the density-dependent cell state model for free vs confined systems. A,** Kymographs of Lgr5 intensity from $t$=0 to $t$=144h, i.e. plotted until day 6. **B**, Radial profiles of Lgr5 and Sca1 for different time-points indicated above each panel column. Predicted intensities are multiplied by the local tissue density.

We next test our model on confinements of different diameters. Independent of parameters, the model does not capture the later reduction of Lgr5 in the center of larger confined systems (diameter 500 – 2000 um) (Fig. M6A, B). Therefore, this model alone cannot capture the spatial patterning of Lgr5 in the system.

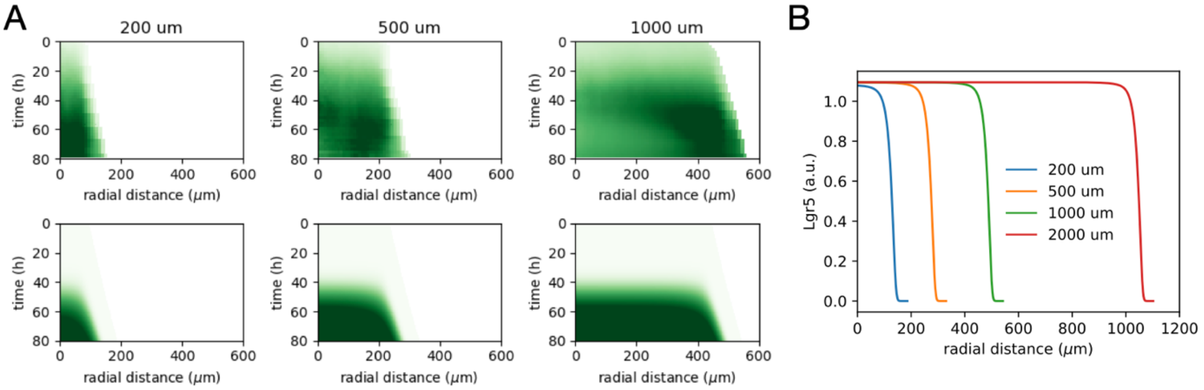


**Fig. M6: Lgr5 dynamics predicted by the density-dependent cell state model for different confinement sizes. A,** Kymographs of Lgr5 intensity in experiment (top row) and model (bottom row). The diameter of the confinement is indicated above each plot. **B**, Spatial pattern of Lgr5 predicted by the model at $t$ = 80h. Predicted intensities are multiplied by the local tissue density.

### 4. Density-dependent translation inhibition model

Based on the finding of density-dependent translation inhibition, we amend the mechano-sensitive model by simulating the dynamics of a generic protein species whose translation is inhibited by compressive stress:

$$\frac{dR}{dt} = I(\rho; K_R, h) - d_R R \tag{17}$$

as a model for the experimental RFP channel, where $I(x; K, h)$ is an inhibiting Hill function

$$I(x; K, h) = \frac{K^h}{x^h + K^h} \tag{18}$$

Since the RFP intensity is already high at the start of the experiment, the initial condition $R(x, t = 0) = 1$ is used. To account for the effect of translation inhibition in the Lgr5 dynamics, we multiply the production term of Lgr5 in Eq. (14) by the translation inhibition function:

$$\frac{dL}{dt} = A(\rho; K_L, h) I(\rho; K_R, h) - d_L L \tag{19}$$

This leads to an optimal density for Lgr5 production (Fig. M7).

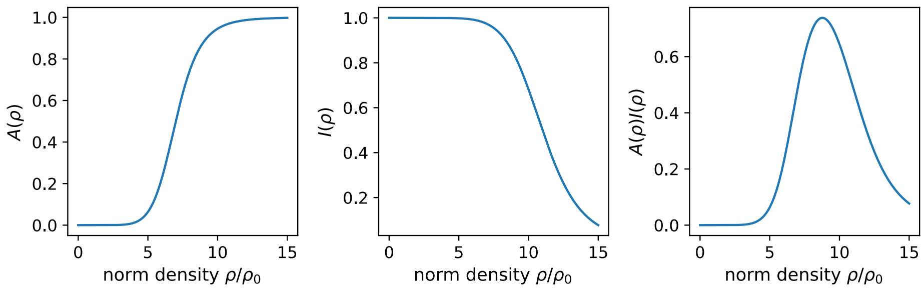


**Fig. M7: Activation and inhibition functions of the density-dependent translation inhibition model.**

Keeping the parameters of the Lgr5 activation unchanged, the parameters of the translation inhibition model (Eq. 16) are fit to recapitulate the RFP and GFP dynamics in the experiment. This model now correctly predicts the spatio-temporal patterning of Lgr5 in the system: initially uniform upregulation, followed by constriction to a ring-shaped Lgr5 positive region for patterns with diameter 500 – 2000 um (Fig. M8A, B). In contrast, in the smaller 200um diameter system, no significant downregulation in the center is predicted, consistent with the experiment. This length-scale emerges due to the pattern of cell density in the system, which increases from the edge inwards with a characteristic length-scale. Due to the optimal cell density in the Lgr5 production model, this translates into a ring-shaped Lgr5 positive region.

The translation inhibition further leads to a reduction in predicted RFP-intensity in the center of the systems (Fig. M8C, D). Thus, the model predicts that the effect of translation inhibition can be removed by normalizing GFP by RFP should lead to patterns as predicted by the original density-dependent Lgr5 model (Fig. M6).

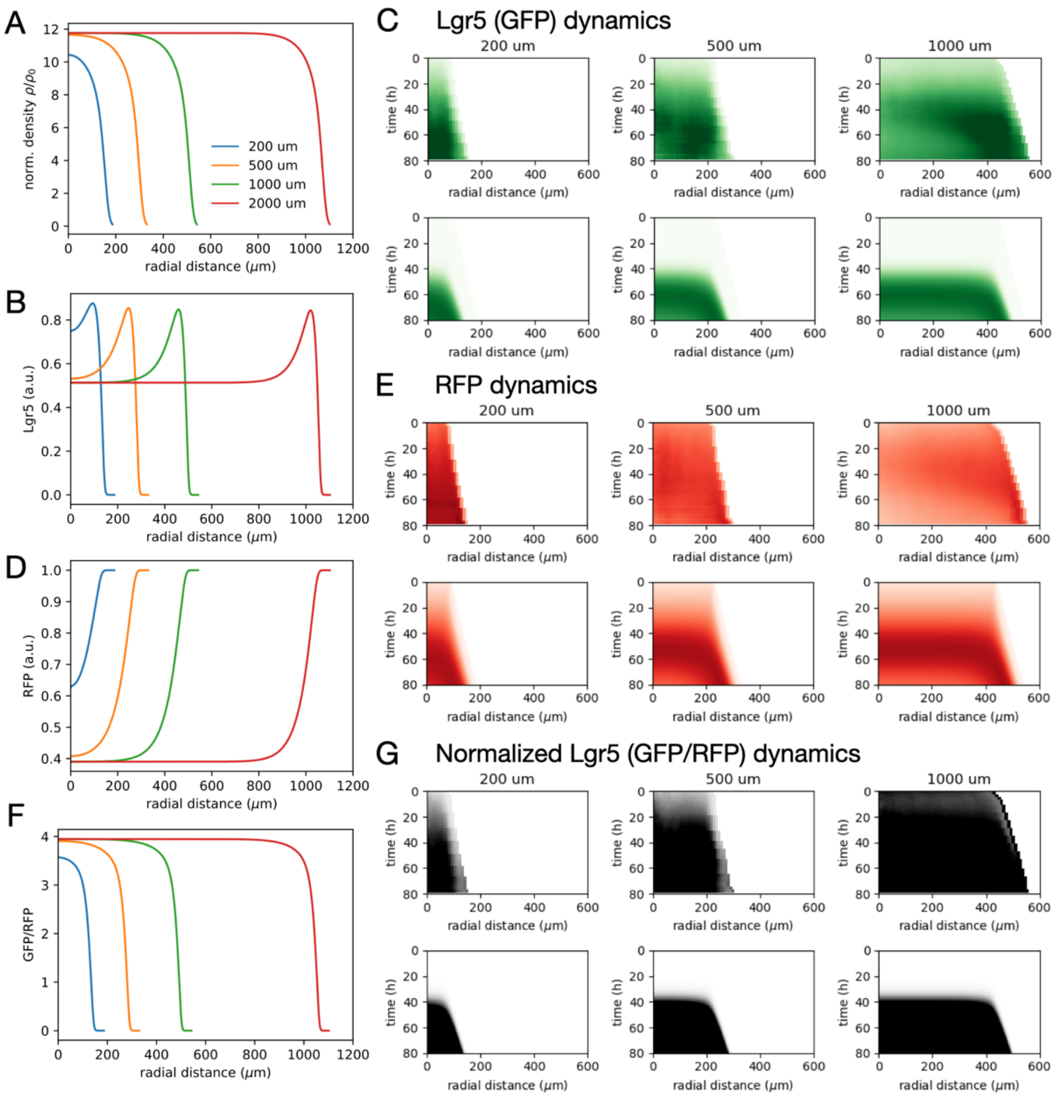


**Fig. M8: Density-dependent translation inhibition Lgr5 model dynamics. A,** Spatial patterns of cell density predicted by the model at $t$ = 80h. **B**, Spatial pattern of Lgr5 predicted by the model at $t$ = 80h. **C,** Kymographs of Lgr5 intensity in experiment (top row) and model (bottom row). **D**, Spatial pattern of RFP predicted by the model at $t$ = 80h. **E**, Kymographs of RFP intensity in experiment (top row) and model (bottom row). **F**, Spatial pattern of GFP/RFP predicted by the model at $t$ = 80h. Predicted intensities are multiplied by the local tissue density. **G,** Kymographs of normalized Lgr5 intensity, GFP/RFP in experiment (top row) and model (bottom row).

### 5. Numerical implementation

The partial differential equations (PDEs) are numerically integrated using custom-written python code using numpy. All simulations are sped up by just-in time-compiling using numba jit.

To numerically integrate Eq. (8), we expand the derivatives:

$$\frac{\partial \rho}{\partial t} = D\left(\frac{\partial^2 \rho}{\partial r^2} + \frac{1}{r}\frac{\partial \rho}{\partial r}\right) - \left(\frac{\rho v_r}{r} + v\frac{\partial \rho}{\partial r} + \rho\frac{\partial v_r}{\partial r}\right) + f(\rho) \tag{18}$$

To ensure numerical stability of Eq. (9), we add a time-derivative of the velocity with a time-scale parameter $\tau = 0.001$ that is chosen to be very small, corresponding to fast equilibration of the system. To ensure physical behaviour, the sign of the time-derivative term is chosen such that the friction term has negative sign.

$$\tau \frac{\partial v_r}{\partial t} = \eta \frac{\partial^2 v_r}{\partial r^2} + \frac{\eta}{r}\frac{\partial v_r}{\partial r} - v_r\left(\zeta + \frac{\eta}{r^2}\right) - \chi \frac{\partial \rho}{\partial r} \quad (19)$$

Then, the time derivative is approximated using Euler forward differences, and the spatial derivative using centered differences. The PDE is simulated on a one-dimensional domain $[0, R_\infty]$. For comparison to experiment, all length-scales are then rescaled by a factor of 100 um, and all time-scales by a factor of 24h.

**6. Parameter overview**

The parameters of the model are summarized below.

| Name | Symbol | Value |
|---|---|---|
| Initial cell density (normalized) | $\rho_0$ | 1 |
| Cell division rate | $k_d$ | 1 |
| Critical cell density | $\rho^*$ | 12.7 |
| Viscosity | $\eta$ | 0.6 |
| Compressibility | $\chi$ | 0.1 |
| Friction, free growth system | $\zeta_{\text{free}}$ | 0.05 |
| Friction, confined system | $\zeta_{\text{confined}}$ | 100 |
| Equilibrium density | $\rho_{eq}$ | 7 |
| Lgr5 degradation rate | $d_L$ | 3 |
| RFP protein degradation rate | $d_R$ | 10 |
| Sca1 degradation rate | $d_S$ | 3 |
| Lgr5 activation threshold | $K_L$ | 6 |
| RFP protein translation inhibition threshold | $K_R$ | 11 |
| Hill exponent | $h$ | 8 |

Table T1: Overview of parameters used in the model, in units with non-dimensionalized time and concentration coordinates.

**Theory references**

### Materials and methods

Organoid culture

Mouse colon tumor organoids (VillinCre-$ER^{T2}$; $APC^{fl/fl}$; $KRAS^{LSL-G12D}$; $TP53^{KO/KO}$ ; R26R-Confetti; $Lgr5^{DTR/eGFP}$) were embedded in Cultrex Basement Membrane Extract (Bio-techne BME001-05) domes and cultured in DMEM/F12 (Gibco) medium supplemented with 100 ng/mL Noggin (CurieCoreTech Recombinant Protein Platform), 1X B27, 1.25 mM N-Acetyl-L-Cysteine and 2X Antibiotic-Antimycotic (Gibco). Organoids were passaged every 3-4 days by mechanical disaggregation.

Mouse tumor models

Animal care and use for this study were performed in accordance with the recommendations of the European Community (2010/63/UE). Experimental procedures were approved by the ethics committee of Institut Curie CEEA-IC #118 (Authorization APAFIS#27460-2020100614277480 v1, APAFIS #38288, and APAFIS#2020101311044955 given by National Authority) in compliance with the international guidelines.

Tumors were generated by injecting trypsinized (TryplE (Gibco) 5 min) single cell suspensions in different organs of NMRI nude 5-6 weeks old female mice (Janvier Labs - Le Genest-Saint-Isle – France). For all procedures, anesthesia was induced using 3% isofluorane (2 L/min airflow) and reduced to 2% isofluorane (0.5 L/min airflow) for the duration of the surgery. At this point, 5 mg/kg Caprofen (Rimadyl) in NaCl 0.9% was injected subcutaneously for analgesia, and tear gel was placed on mice eyes to prevent drying. Mice were maintained at 37°C for the duration of anaesthesia using heating pads. The specific protocols for each tumor model were as follows:

- Interscapular fat pad: an incision was opened at the skin of the interscapular region and the fat pad was pinched out using tweezers. 100 µL of organoid culture media containing 500.000 cells was injected into the fat pad using a 25G needle. The incision was sutured using clips.
- Colon: colon orthotopic tumors were generated as described previously(*1*). Briefly, the colon mucosa was damaged using an electric interdental brush (Panasonic EW0945) previously soaked in a 0.5M EDTA solution at 50°C and pH8. 100 µL of a 1:1 matrigel-cell solution containing 500.000 cells were injected in the colon using a p200 pipette and the anus was closed using surgical glue to avoid leaking of the cell suspension.
- Liver: a lateral skin incision was opened, and the spleen was exteriorized using tweezers. 100 µL of organoid culture media containing 500.000 cells were injected in the tip of the spleen using a 25G needle. The spleen was reinserted in the mouse and the incision was sutured using clips.
- Subcutaneous: for external compression experiments, culture media containing 356 000 cells was injected subcutaneously in the interscapular region of NMRI nude mice.

Organoid monolayer culture

Organoid monolayers were generated from trypsinized single-cell suspensions obtained from 3D organoid cultures. Briefly, 3D organoids were isolated from BME and mechanically disaggregated using an 18G needle. Cell suspension was centrifuged at 300 RCF for 3 minutes, resuspended in 1 mL of TryplE and incubated at 37°C for 5 minutes. Cells were then mechanically disaggregated by vigorously pipetting up and down and TryplE was inactivated by adding 9 mL of PBS containing 10% FBS. Cells were again centrifuged at 400 RCF for 5 minutes and resuspended in experiment medium: DMEM-F12, 1.25 mM N-Acetyl-L-Cysteine, 1X B27, 2X Antibiotic-Antimycotic, and

Metronidazole/Ciprofloxacin (20 and 5 μg/mL, respectively). 50 μL of cell solution containing ~200.000 cells were seeded on collagen-coated polyacrylamide gels or filters.

- Polyacrylamide gel fabrication and functionalization: collagen-coated 5kPa (Young's modulus) polyacrylamide gels were fabricated as previously described (*2*). Briefly, glass-bottom dishes (World Precision Instruments) were treated for 15 min with a solution of silane (3-(Trimethoxysilyl) propyl methacrylate, Sigma-Aldrich) diluted 1:3 in PBS. After 3 washes with water (5 minutes each), dishes were incubated with a solution of 0.5% Glutaraldehyde/PBS (w/v) for 30 min. After 3 more washes with water (5 minutes each) and through drying, 5kPa polyacrylamide gels were fabricated by placing a 20 μL drop of polyacrylamide solution on the glass bottom, covered by an 18 mm coverslip. The polyacrylamide solution contained 382.95 mL of PBS, 93.3 μL of 40% Acrylamide, 11 μL of 2% Bis-acrylamide, 10 μL of 0.2 μm carboxylate-modified microspheres (ThermoFisher Scientific), 2.5 μL of 10% ammonium persulfate (VWR) and 0.25 μL of N, N, N', N'-tetramethylethylenediamine (Sigma-Aldrich). For traction force microscopy, micropheres were fluorescent (F8807, ThermoFisher Scientific), for immunostaining, non-fluorescent microspheres were used (C37480, ThermoFisher Scientific). After 1 hour of polymerization at room temperature, the coverslip was removed, and the gel was incubated in PBS. For functionalization, gels were incubated with 75 μL of a 2 mg/mL Sulpho-SANPAH (Sigma-Aldrich) solution under UV light (365nm) for 7.5 minutes. This solution was washed twice with 10 mM HEPES and once in PBS (3 min each), followed by gel drying for 5 minutes. At this point, a 10 μg/mL solution of rat tail collagen I (Corning) was added on top of the gel and incubated overnight at 4°C.
- Patterning: collagen and organoid patterning were performed as previously described (*3*). Briefly, polydimethylsiloxane (PDMS) stencils containing openings with the desire size and shape were fabricated by spinning PDMS on Su8 masters generated by soft lithography. The stencils were incubated with a solution of pluronic acid F127 (Sigma-Aldrich) 2% in PBS for 1 hour, washed twice with PBS and let dry for 20 minutes. Stencils were then carefully placed on top of polyacrylamide gels at different steps of gel fabrication depending on the experiment. To pattern the initial organoid shape in free growth experiments (Fig. 1 and 2), stencils were added on collagen I coated gels right before organoid seeding. To generate collagen I patterns and grow organoids in a confined environment (Fig 2, 4-6), stencils were placed on Sulpho-SANPAH treated polyacrylamide gels right before adding the Collagen I solution. Collagen I patterns of different radii were combined in the same gel for live imaging.
- Filter functionalization: 0.4 μm Transwell filters (Corning) were coated with neutralized rat tail collagen-1 solution (2mg/ml; Corning). 200-400 μL of solution was spread as much as possible on the filter using a 200 μL pipette and the solution was let polymerize at 37°C with humidity control for 1 hour before organoid seeding.

Immunostaining

- Tumors: All tumors except for compressed tumors (Fig. 2G-I) were washed in PBS and fixed in 4% paraformaldehyde (PFA) during 45 minutes for small tumors and 2 hours for large tumors, shaking at RT. After washing with PBS for 10 min, tumors were incubated with 15% sucrose in PBS shaking for 2h, and later transferred to 30% sucrose in PBS overnight at 4°C. Tumors were embedded in OCT (Sakura) and fully sectioned in sequential 10 μm thick slices attached to SuperFrost Plus™ Adhesion slides (VWR, Menzel Gläser). Immunostainings were performed at equatorial tumor sections. Sections were permeabilized with 0.2% Triton X-100 in PBS for 1h at

RT, blocked with 3% BSA in PBS containing 0.05% Triton X-100 for 1h at RT and incubated with primary antibodies overnight at RT. Sections were then washed 3 times (10 min each) and incubated with secondary antibodies and DAPI for 4h at RT. After 3 more washes (10 min each), sections were mounted using AquaPolyMount (Polysciences). All washes were done in 0.05% Triton X-100 in PBS, and antibody incubations were done in blocking solution.

Immunostainings of compressed tumors (Fig. 2G-I) were performed using the OPAL protocol (Akoya Biosciences) on the Leica BOND RXn autostainer (Leica Microsystems).

- Organoids: organoids were fixed in 4% PFA for 20 min at RT and washed 3 times (5 min each) with PBS. Samples were permeabilized with 0.5% Triton X-100 in PBS for 30 min at RT, blocked with 10% FBS in PBS for 1h at RT and incubated with primary antibodies overnight at 4°C. Samples were then washed 3 times (5 min each) and incubated with secondary antibodies and DAPI for 1h at RT. After 3 more washes (10 min each), samples were mounted using AquaPolyMount (Polysciences). All washes were done in PBS, and antibody incubations were done in blocking solution.

Antibodies

Primary antibodies: Sca1 (ab51317, Abcam, 1:500), Ki67 (ab15580, Abcam, 1:200), CC3 (9661, Cell Signaling Technology, 1:200), GFP (A-11122, Thermo Fisher, 1:250) with Opal 520, RFP (600-401-379, Rockland, 1:250) with Opal 690. Secondary antibodies: Donkey anti-Rat Alexa Fluor 647 (A-78947, Thermo Fisher, 1:400), Donkey anti-Rabbit 647 (A-31573, Thermo Fisher, 1:400).

EdU and OPP pulses

Organoids were incubated with medium containing EdU (10 µM, Thermo Fisher Scientific) or OPP (20 µM, Thermo Fisher Scientific) for 30 min before fixation in 4% PFA for 20 min at RT. After 3 washes, organoids were permeabilized with 0.5% Triton X-100 in PBS for 30 min at RT and click-it reaction for EdU or OPP revelation was performed (according to the manufacturers protocol) using the Alexa Fluor 647 dye. After 3 washings (5 min each), samples were incubated with a 1:200 DAPI solution for 15 min, washed and mounted using AquaPolyMount (Polysciences).

Imaging

- Tumors: tumor slices were imaged using a AXR (Nikon) or an LSM900 (Zeiss) scanning confocal microscope. Full tumor slices were imaged using a 10x objective (Air, NA 0.45) and state-density relations were measured using a 60x objective (Oil, NA 1.42). Whole-mount imaging for stress measurements in tumors was performed on a W1 spinning disk (Yokogawa) using a 40x objective (WI, NA 1.25).
- Organoids: immunostainings and live imaging were performed on a W1 spinning disk (Yokogawa) using a 40x (WI, NA 1.25) and 20x (Air, NA 0.45) objectives, respectively. Microscopes were equipped with temperature, $CO_2$ and humidity control.

Tumor stress measurements

Polyacrylamide beads were fabricated and characterized as previously described (*4*, *5*). Briefly, a droplet-based microfluidic platform with flow-focusing geometry was employed to emulsify a polyacrylamide pre-gel solution in fluorinated oil (3M™ Novotec™ 7500, Iolitec Ionic Liquids Technologies GmbH, Germany). The polyacrylamide pre-gel solution contained 40% acrylamide (Sigma-Aldrich) monomer, 2% bis-acrylamide (Sigma-Aldrich) crosslinker and 0.05% ammonium persulfate (Sigma-Aldrich) as the radical initiator, prepared in 10 mM Tris buffer (pH 7.5). The fluorinated oil contained 2% ammonium-Krytox® surfactant and 0.4% N,N,N′,N′-

tetramethylethylenediamine (Sigma-Aldrich) as a catalyst and 0.1% acrylic acid N-hydroxysuccinimide-acrylamide ester (Sigma-Aldrich) to enable covalent coupling of fluorophores and proteins. The beads were functionalized with Alexa Fluor™ 633 (ThermoFisher Scientific) and poly-D-lysine (Gibco) to allow fluorescence detection and cell adhesion. Following polymerization, the beads were washed, suspended in 1×PBS and stored at 4 °C. Bead size distributions were quantified from bright-field images using a custom Fiji macro and the Young's modulus of individual beads was determined by atomic force microscopy (AFM). 500.000 beads were mixed with 500.000 cancer cells and injected into the interscapular fat pad of NMRI nude mice as described above. Tumors were harvested for 1 or 2 weeks, fixed in PFA 4% for 2h, washed, permeabilized for 24h in PBS containing 2% Triton X-100 and incubated in RapiClear 1.49 (SunJin Lab) for at least 3 days shaking at 4°C.

Tumor confinement

When subcutaneous tumors reached around 100 mm3 (± 20 mm3), the compressive device (*6*) (holder+screw) was implanted using stitches and maintained for 4 days (Fig. S7A,B,D). The imposed pressure was calibrated using a force sensor (FlexiForce® OEM Kit, Teckscan, South Boston, MA, USA) inserted between the subcutaneous tumor and the screw. Every day, the stitches were checked and restored if removed from the mouse. After 4 days, compressed tumor volume was calculated and compared to uncompressed tumors (Fig. S7C). Tumors were then harvested and fixed for 24 hours in PFA 4%, dehydrated in xylene/methanol and embedded in paraffin according to IUCT-Oncopole protocol. Paraffin was sectioned in 7 μm thick slices and used for immunostaining.

Image analysis

Image analysis was performed using custom-made software programmed in ImageJ and Matlab.

- Image stitching: stitching was performed by pre-aligning the images based on microscope stage positions and drift was calculated using 2D cross-correlation. Upon aligning the images, a linear blending of a user-defined length was applied in overlapping regions before combining all the images into a final stitched image. For traction force microscopy, alignment was performed using reference bead images after organoid trypsinization.
- Particle-Image Velocimetry (PIV): PIV was performed on the RFP channel as described previously (*2*). Briefly, local velocities were measured by calculating the cross-correlation of overlapping (75%) imaging windows (64x64 pix) between consecutive timepoints (2h timestep) using a custom-made software programmed in Matlab.
- Traction Force Microscopy (TFM): 2D TFM was performed as described previously (*7*). Images of fluorescent beads were taken during the timelapse imaging. Bead displacement was calculated by performing PIV of every bead image compared to a reference bead image were organoids had been removed by trypsinization. 2D traction force fields were calculated from displacements using finite-thickness Fourier transform TFM using a custom-made software programmed in Matlab.
- Monolayer Stress Microscopy (MSM): MSM was performed as described previously (*8*). Briefly, force balance with tractions yields the tension field in the monolayer, as a second rank symmetric tensor. Average principal stress was used for quantifications.
- Quantification of radial profiles: tumor cells and organoids were automatically segmented by applying a gaussian filter and an intensity-threshold on RFP images. This segmentation was manually corrected when errors were detected. The resulting cell mask was used to compute the closest linear distance of each point of the tissue to the edge. The magnitude of interest (fluorescent intensity, cell density, velocity, traction force and intercellular stress) was averaged according to its distance to the edge. Vectorial variables, such as traction and velocity, were previously

decomposed in radial and tangential components in respect to the closest tissue edge, as previously described (*2*). Different replicates were resized to the mean length within each condition before averaging. For kymographs, resizing was performed at every timepoint. For fluorescent intensity, background noise was subtracted from the image before any quantification.

- Quantification of cell density: When possible, cell nuclei were automatically segmented using Cell Pose (*in vitro*, free growth up to day 4 and confined growth at day 0). For all other conditions, *in vitro* and *in vivo*, nuclei were manually counted using multipoint selection in ImageJ. Nuclear coordinates were imported to Matlab and local cell density was calculated using a moving window of 100 x 100 µm. In each window, average Lgr5 EGFP and Sca1 intensity within the tissue (RFP+ regions) was also quantified. All windows from all conditions were combined to plot state-density relations. Density maps were also used to compute radial density profiles.
- Lgr5 autocorrelation distance: this analysis was performed on central regions of 3-weeks tumors exhibiting Lgr5 periodic pattern. After applying a gaussian filter to the Lgr5 channel, 2D autocorrelation was calculated and the mean minimum cartesian distance between all the autocorrelation peaks was computed. Peaks were detected using peaks2 function in Matlab (Kristupas Tikuišis (2025).
- Linescans of hierarchichal units: hierarchical units were visually identified based on Lgr5 EGFP signal and manual linescans (50 pixels thickness, ImageJ) were performed to calculate fluorescence intensity profiles. Average Lgr5 EGFP, RFP, Sca1, Ki67 and Cleaved-caspase 3 along the line were calculated for each hierarchical unit. To average linescans, each independent measurement was aligned according to the Lgr5 EGFP central intensity peak defined by fitting a gaussian function.

Single-cell RNA sequencing

- Cell isolation: Stencils containing grids of 200 µm or 1000 µm circles were fabricated. Both grids contained a comparable patterned area to minimize differences in cell numbers between conditions. Organoids were seeded on patterns every 2 days and all conditions (Day 0, 2 and 4) were processed at the same time to minimize badge effects. Cell isolation was performed by trypsinization (TryplE) at 37 °C for 10 min, mechanical disaggregation by pipetting and filtering through a cell strainer (40 µm). 5000 cells per condition were processed through the 10x genomics single cell sequencing platform and sequenced at 100.000 reads per cell.
- scRNA-seq pre-processing: single-cell data analysis was performed by GenoSplice technology (www.genosplice.com). Sequencing data quality was assessed using FastQC v0.11.5. For read alignment and unique molecular identifiers (UMI) quantification, the CellRanger software v7.1.0 was used on Mouse reference data 2020-A (genome mm10, gene annotation ensemble) with default parameters. In order to estimate and suppress ambient RNA, DecontX [PMID:32138770] R package was applied on counts data from cellranger. The 2 expression matrices containing the UMI counts were merged, and only the genes with UMI ≥ 1 in at least one cell were kept. The following filters were applied to generate a global matrix used in further analysis: number of detected genes ≥ 3000, and cells with UMI in mitochondrial genes ≤ 15%. DoubletFinder [PMID:30954475] was used to suppress doublet. Finally, an integration step using Harmony [PMID:31740819] was performed. The resulting sample after the pre-processing was composed of 14363 single cell transcriptomes from all conditions.
- Cell cycle: According to the marker genes of cell cycle, the cell cycle of each cell was counted using Seurat package. Then, all cells were divided into G1, G2M and S phases according based on their scores.

- Clustering: For normalization and clustering, Seurat 5.2.1 was used [PMID:29608179,34062119], and SCTransform normalization was applied. Based on elbowplot, 31 PC were used for UMAP calculation and clustering analysis. Clustering step was performed using default parameters from Seurat on haromany reduction (FindNeighbors and FindClusters functions). To calculate markers for each cluster, a global-scaling normalization method was applied with a scale factor of 10,000 and log-transformation of data. Only genes expressed in at least 15% of cells with a log2FC minimum of 0.25 a adjusted p-value inferior at 0.05 were considered as markers using Seurat Wilcoxon test. Based on treemap of clusters with different resolution parameters and clusters markers, we choose a resolution parameter of 0.4. For signature analysis, AddModuleScore from Seurat package was used.
- Differential expression analysis: Analysis for enriched GO terms, KEGG pathways and REACTOME pathways were performed using WebGestaltR package [PMID: 28472511], on databases from Human organism. GO terms and pathways were considered as enriched if fold enrichment ≥ 2.0, uncorrected p-value ≤ 0.05 and minimum number of regulated genes in pathway/term ≥ 2.0.
- Figures: For graphics representation, functions from Seurat (VlnPlot, FeaturePlot …) and scCustomize package were used.

**Methods references**


1. S. Richon, O. Zajac, C. Perez Gonzalez, D. Matic Vignjevic, Optimized protocol for the generation of an orthotopic colon cancer mouse model and metastasis. *STAR Protoc* **4**, 102022 (2023).
2. C. Pérez-González, G. Ceada, F. Greco, M. Matejčić, M. Gómez-González, N. Castro, A. Menendez, S. Kale, D. Krndija, A. G. Clark, V. R. Gannavarapu, A. Álvarez-Varela, P. Roca-Cusachs, E. Batlle, D. M. Vignjevic, M. Arroyo, X. Trepat, Mechanical compartmentalization of the intestinal organoid enables crypt folding and collective cell migration. *Nat Cell Biol*, doi: 10.1038/s41556-021-00699-6 (2021).
3. C. Pérez-González, R. Alert, C. Blanch-Mercader, M. Gómez-González, T. Kolodziej, E. Bazellieres, J. Casademunt, X. Trepat, Active wetting of epithelial tissues. *Nat Phys* **15**, 79–88 (2019).
4. R. Goswami, K. Kim, A. R. Boccaccini, J. Guck, S. Girardo, Fine-tuning cell-mimicking polyacrylamide microgels: Sensitivity to microscale reaction conditions in droplet microfluidics. *bioRxiv*, 2025.07.24.666603 (2025).
5. N. Träber, K. Uhlmann, S. Girardo, G. Kesavan, K. Wagner, J. Friedrichs, R. Goswami, K. Bai, M. Brand, C. Werner, D. Balzani, J. Guck, Polyacrylamide Bead Sensors for in vivo Quantification of Cell-Scale Stress in Zebrafish Development. *Sci Rep* **9**, 1–14 (2019).
6. M. Di-Luoffo, S. Arcucci, N. Therville, T. Marty, R. D'Angelo, M. Chaouki, B. Thibault, P. Swinder, P. Assemat, M. Delarue, J. Guillermet-Guibert, Mutants of p53 sustains tumor growth under mechanical compression. *bioRxiv* (2025).
7. X. Trepat, M. R. Wasserman, T. E. Angelini, E. Millet, D. A. Weitz, J. P. Butler, J. J. Fredberg, Physical forces during collective cell migration. *Nat Phys* **5**, 426–430 (2009).
8. D. T. Tambe, C. C. Hardin, T. E. Angelini, K. Rajendran, C. Y. Park, X. Serra-picamal, E. H. Zhou, M. H. Zaman, J. P. Butler, D. A. Weitz, J. J. Fredberg, X. Trepat, Collective cell guidance by cooperative intercellular forces. *Nat Mater* **10**, 469–475 (2011).
